# *De Novo* Design of Orthogonal Far-Red, Orange, and Green Fluorophore-binding Proteins for Multiplexed Imaging

**DOI:** 10.1101/2025.08.03.668343

**Authors:** Long Tran, Steffen Klein, David Juergens, Shajesh Sharma, Justin Decarreau, Gyu Rie Lee, Yujia Wang, Wei Chen, Asim K. Bera, Alex Kang, Jon Woods, Emily Joyce, Dionne K Vafeados, Nicole Roullier, Xinting Li, Bingxu Liu, Yang Bo, Edin Muratspahić, Tim A. Brown, Jonathan B. Grimm, Ronak Patel, Luke D. Lavis, Julia Mahamid, Linna An, David Baker

**Affiliations:** Department of Chemical Engineering, University of Washington, Seattle, WA, USA; Institute for Protein Design, University of Washington, Seattle, WA, USA; Molecular Systems Biology Unit, European Molecular Biology Laboratory (EMBL), Heidelberg, Germany; Department of Biochemistry, University of Washington, Seattle, WA, USA; Graduate Program in Molecular Engineering, University of Washington, Seattle, WA, USA; Department of Chemistry, Stanford University, Stanford, CA, USA; Department of Bioengineering, University of Washington, Seattle, WA, USA; Department of Biological Sciences, Korea Advanced Institute of Science and Technology, Daejeon, South Korea; Janelia Research Campus, Howard Hughes Medical Institute, Ashburn, Virginia, USA; Cell Biology and Biophysics Unit, European Molecular Biology Laboratory (EMBL), Heidelberg, Germany; Howard Hughes Medical Institute, University of Washington, Seattle, WA, USA

**Author notes:** These authors contributed equally as first authors (Long Tran, Steffen Klein). These authors contributed equally as second authors (David Juergens, Shajesh Sharma, Justin Decarreau).

## Abstract

Fluorescent proteins and small-molecule dyes offer complementary advantages for biological imaging: proteins are amenable to genetic tagging, whereas dyes provide superior brightness and photostability. To combine these strengths, we used *de novo* protein design to generate small, nanomolar-affinity, high-selectivity binders (NovoTags) for three cell-permeable dyes spanning the visible spectrum. We show that the NovoTag fluorescent lifetimes can be tuned, and demonstrate their application in lifetime and wavelength-based multiplexed fluorescence imaging. We further design a two-chain NovoTag that functions as a chemically induced dimerization system with fluorescent readout in living cells, or as a minimally perturbing proximity probe in fixed cells. Our approach combines the advantages of fluorescent proteins and small-molecule dyes, expanding the toolkit for cellular imaging.

## Introduction

Biological imaging has been transformed by the ability to tag specific cellular components with genetically encoded fluorescent proteins, but fluorescent proteins have limited brightness and photostability, constraining the resolution, duration, and complexity of imaging experiments (*1*). In contrast, small-molecule dyes offer superior optical properties (*2–4*); advanced synthetic fluorophores, such as the Janelia Fluor (JF) dyes, are notable for their quantum yields across a range of emission wavelengths, photostability, and cell permeability (*2*, *3*). However, the use of JF dyes in cellular imaging requires a targeting mechanism to the proteins of interest. Current approaches primarily employ fusion tags, including HaloTag (*5*) and SNAP-tag (*6*), which covalently bind modified versions of the dyes. While effective, these tags are relatively large (20-35 kDa) and not dye-specific; they bind a general ligand moiety that can be conjugated to a compatible dye (*5*, *6*). This has complicated the development of orthogonal versions, thereby limiting multiplexing capabilities (*7*).

We reasoned that designing a set of small proteins, each selectively binding to a specific JF dye across the visible spectrum, could provide a general solution to the multiplexed imaging problem. This, however, poses a challenging design task because JF dyes are structurally analogous rhodamine derivatives that lack known natural binders (*8*). To address this challenge, we set out to leverage recent advances in machine-learning-based *de novo* protein design (*9*, *10*) to generate genetically encodable, small, JF dye-binding proteins with high affinity and specificity for single dyes, and to explore the use of such designs for multiplexed fluorescence imaging.

## Results

### Design and evaluation of fluorophore-binding proteins

We selected three JF dyes (JF_494_, JF_596_, and JF_657_) (**Fig. 1A – C**) with well-separated excitation and emission spectra spanning the visible range (*2*) as design targets (**Fig. 1D, Fig. S1**). These fluorophores share a common rhodamine-derived core structure but differ in specific chemical modifications that enhance brightness and tune their excitation and emission maxima (*2*). We reasoned that a design strategy that generates proteins with extensive complementarity to an input ligand structure could achieve specificity, as even small changes in the chemistry could introduce clashes or eliminate favorable contacts (*9*, *11*).

**Fig. 1.**
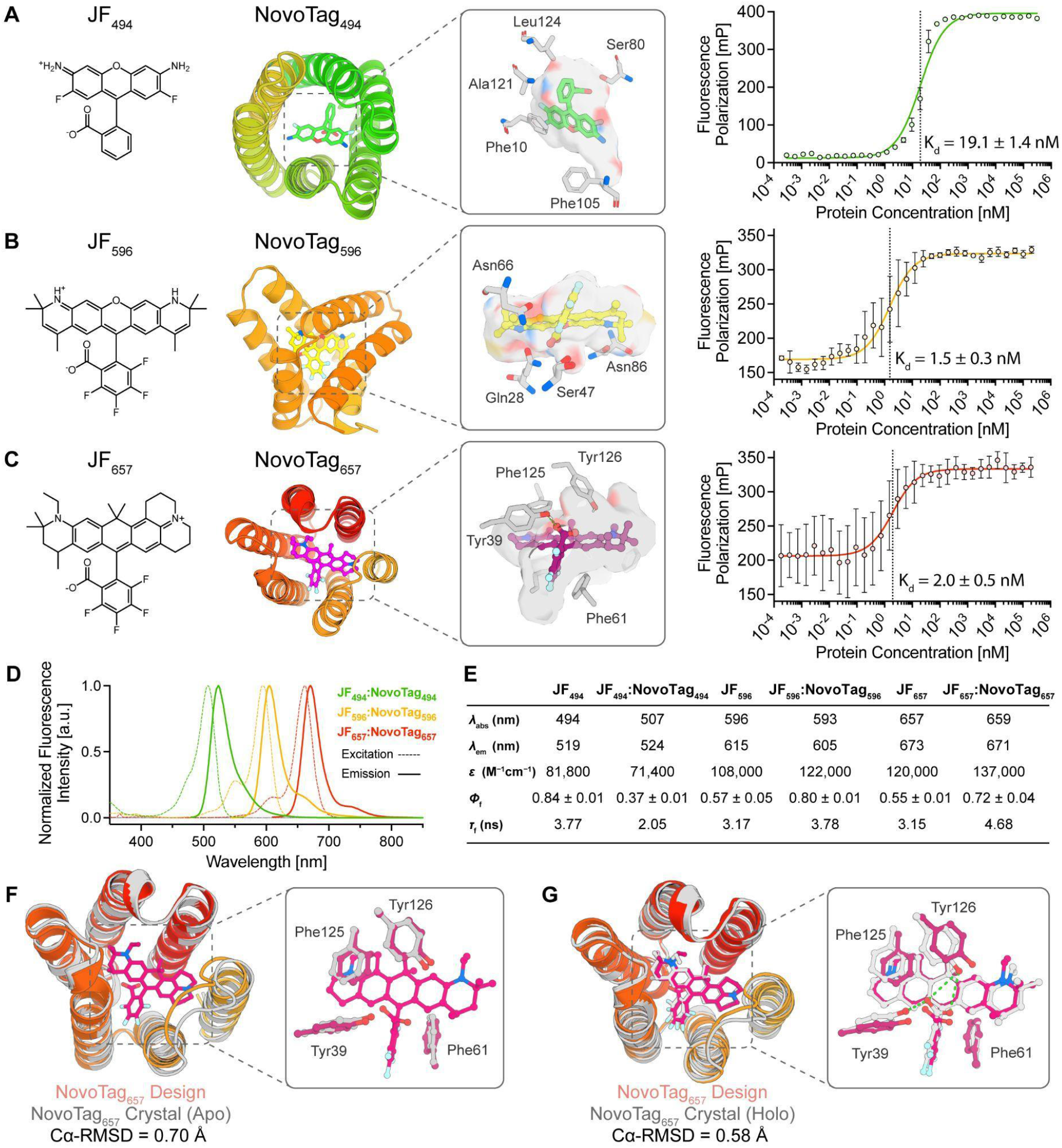
Design and characterization of Janelia Fluor-binding proteins. **(A – C)** NovoTag designs for the binding of JF_494_ (A), JF_596_ (B), and JF_657_ (C). For each JF dye–NovoTag pair, the chemical structure of the target dye (left), model of the NovoTag design (middle) with the relevant dye-interacting residues highlighted (insets), and fluorescence polarization (FP) titration curves (right) are shown. For each FP titration, 3-8 independent measurements were acquired. For each concentration, the FP (mean ± SD) is shown. A non-linear regression (standard binding isotherm model) was fitted, and K_d_ values were determined and reported with standard error (SE). **(D)** Excitation (dotted lines) and emission (solid lines) of JF dye-NovoTag pairs: JF_494_:NovoTag_494_ (green), JF_596_:NovoTag_596_ (orange), and JF_657_:NovoTag_657_ (red). **(E)** Photophysical characterization of free JF dyes and JF dye–NovoTag conjugates: Absorbance maximum (λ_abs_), emission maximum (λ_em_), extinction coefficient (ε), fluorescence quantum yield (Φf) with SD, and fluorescence lifetime (*τ*_f_). **(F & G)** Superposition of the design model of NovoTag_657_ (red) with the apo (F) and holo (G) crystal structure (gray). Green dashed lines indicate hydrogen bonds.

We developed a machine learning based approach for generating binders with internally repeating closed structures (pseudocycles) that encircle the target JF dyes (pseudocyclic protein scaffolds have been previously found to be advantageous for small molecule binding, but have not been implemented in the ligand-aware context (*9*)). We used the generative deep learning method Cα RFdiffusion (*10*) (**Fig. S2**) with symmetry conditioning to build protein structures around the JF_494_, JF_596_, and JF_657_ dyes. We encoded the ligand structure via the Cα RFdiffusion motif template input and omitted symmetry operations in any ligand-ligand or ligand-protein blocks in the pair representation (Methods). We found that this method effectively generated monomeric pseudocyclic structures around the three selected JF dyes (**Fig. 1A – C**). This RFdiffusion-based method has the advantage over our previous hallucination-based approach (*9*) as it generates pseudocyclic proteins directly around a target ligand, thus avoiding a second ligand-docking step. After obtaining the structures, we generated sequences using LigandMPNN, with FastRelax cycling (*12*). Each design comprised a single-domain protein of 110-160 amino acids, with a predicted cavity tailored to accommodate the targeted JF dye. We filtered the designs with Rosetta (*13*) for ligand binding and with AlphaFold2 (AF2) (*14*) for protein sequence-structure consistency. A total of 4843, 5800, and 6032 binder variants were selected for JF_494_, JF_596_, and JF_657_ binding, respectively. Those were synthesized on oligonucleotide arrays, displayed on the yeast surface, and screened by fluorescence-activated cell sorting (FACS) (**Fig. S3**). Next-generation sequencing of enriched pools revealed multiple high-affinity binders: 56 designs for JF_494_, 236 designs for JF_596_, and 334 designs for JF_657_ exhibited binding affinities on yeast of 5 µM or lower.

For each dye, we selected a single binder design based on the FACS enrichment followed by single clone flow cytometry signal intensity for subsequent characterization (**Fig. S3**). Those were expressed in *E. coli*, purified by affinity chromatography, and binding affinities were measured using fluorescence polarization (FP) assays. The selected designs for each target JF dye (NovoTag_494_, NovoTag_596_, and NovoTag_657_ for JF_494_, JF_596_, JF_657_) have molecular weights of 15.1 kDa, 13.3 kDa, and 15.6 kDa, and binding affinities of K_d_ = 19.1 nM, K_d_ = 1.5 nM, and K_d_ = 2.0 nM, respectively (**Fig. 1A – C**). We used stopped-flow fluorescence polarization to determine the binding kinetics of JF_494_ to NovoTag_494_, yielding a k_on_ of 1.8 × 10^5^ M^-1^s^-1^ and k_off_ of 0.00336 s^−1^ (**Fig. S4**), consistent with the measured binding affinity of K_d_ = 19.1 nM (**Fig. 1A**). Next, we assessed the specificity of the NovoTags to their JF targets using FP and found that they bound their respective targets with at least 1,000-fold greater affinity than the alternative ligands (**Fig. S5**). We further characterized the absorbance maximum (*λ*_abs_), emission maximum (*λ*_em_), extinction coefficient (*ε*), fluorescence quantum yield (*Φ*_f_), and fluorescence lifetime (*τ*_f_) of the free dyes and their conjugates (**Fig. 1E**). For both NovoTag_657_ and NovoTag_596_, we observed modest shifts in *λ*_abs_ and *λ*_em_, and larger *ε* (∼ 15%) and *Φ*_f_ (∼ 40%) values upon protein binding, resulting in a substantial increase in fluorescence emission intensity of the protein–dye complexes compared to the free dyes. In contrast, NovoTag_494_ showed a decrease in both *ε* (∼15%) and *Φ*_f_ (∼50%). The measured *τ*_f_ values matched the trends in *Φ*_f_, with NovoTag_657_ and NovoTag_596_ increasing the fluorescence lifetime and NovoTag_494_ decreasing it compared to the free dyes. Compared to the state-of-the-art synthetic fluorophore JF_646_-HaloTag ligand bound to HaloTag7 (ε = 152,000 M^-1^cm^-1^, *Φ*_f_ = 0.54) (*3*) or JF_657_-HaloTag ligand bound to HaloTag7 (ε = 128,000 M^-1^cm^-1^, Φ = 0.54), JF_657_ bound to NovoTag_657_ exhibited 20% and 43% higher brightness, respectively (ε = 137,000 M^-1^cm^-1^, *Φ*_f_ = 0.72).

We characterized the biophysical properties of the NovoTag designs using circular dichroism (CD) and size exclusion chromatography (SEC). CD spectra revealed predominantly α-helical secondary structure consistent with our computational models, and thermal denaturation experiments indicated high thermostability (**Fig. S6**). SEC traces showed that NovoTags are highly soluble and monomeric in solution.

We determined the crystal structures of NovoTag_657_ in the apo and holo states at 1.44 Å and 2.37 Å resolution, respectively, showing that they closely match the design model over the protein backbone (Cα-RMSD-design-holo = 0.58 Å, Cα-RMSD-design-apo = 0.70 Å, **Fig. 1F & G, Table S1, S2**). The aromatic side chains of Phe61 and Phe125, designed to form hydrophobic contacts and π-π stacking interactions with the dye, as well as Tyr39 and Tyr126, which form hydrogen bonds with JF_657_, were correctly positioned. All-by-all AlphaFold 3 (AF3) (*15*) prediction comparisons of NovoTags in complex with JF dyes yielded highest confidence predictions (highest ipTM, highest ligand pLDDT) for the designed on-target pairs (**Fig. S7)**. These structures and in silico results show how the high protein-small molecule shape complementarity generated by our design methods can lead to specificity even among small-molecule analogs with high structure similarity (**Fig. S5**)

### NovoTags enable multiplexed fluorescence microscopy in cells

We next evaluated each NovoTag as a genetically encoded fluorescent tag for microscopy. We fused the NovoTags either to mScarlet (NovoTag_494_) or eGFP (NovoTag_596_ and NovoTag_657_) and targeted the constructs to the outer mitochondrial membrane using a mitochondrial localization signal derived from Tom70 (*16*) (MitoTag) in HeLa cells. Cells were incubated with the corresponding JF dye (5 nM for 15 min). Following extensive washing, live-cell fluorescence imaging showed that the JF dye signal colocalized with the mScarlet or eGFP signal on the mitochondrial membrane (**Fig. S8**), demonstrating that all three NovoTags effectively bind and recruit the cell-permeable JF dyes to specific subcellular sites. The nanomolar affinities (**Fig. 1A – C**) and slow k_off_ rate (**Fig. S4**) of the NovoTags enabled live-cell imaging after washing, thereby minimizing nonspecific background, highlighting the benefits of high-affinity binders. To assess the impact of chemical fixation on NovoTag staining, MitoTag-NovoTag_657_-eGFP was expressed in HeLa cells, and JF dye staining was performed either before or after fixation (**Fig. S9**). In both cases, the JF_657_ signal colocalized with eGFP, demonstrating that the NovoTag system is compatible with this common imaging workflow. Comparing the photostability of NovoTag_657_ with that of the well-established, commercially available JFX_650_-HaloTag ligand (*17*) showed comparable rates of photobleaching in HeLa cells with 50% signal loss after 14 or 15 bleaching iterations, respectively (**Fig. 2C, Fig. S10, Movie S1**).

**Fig. 2.**
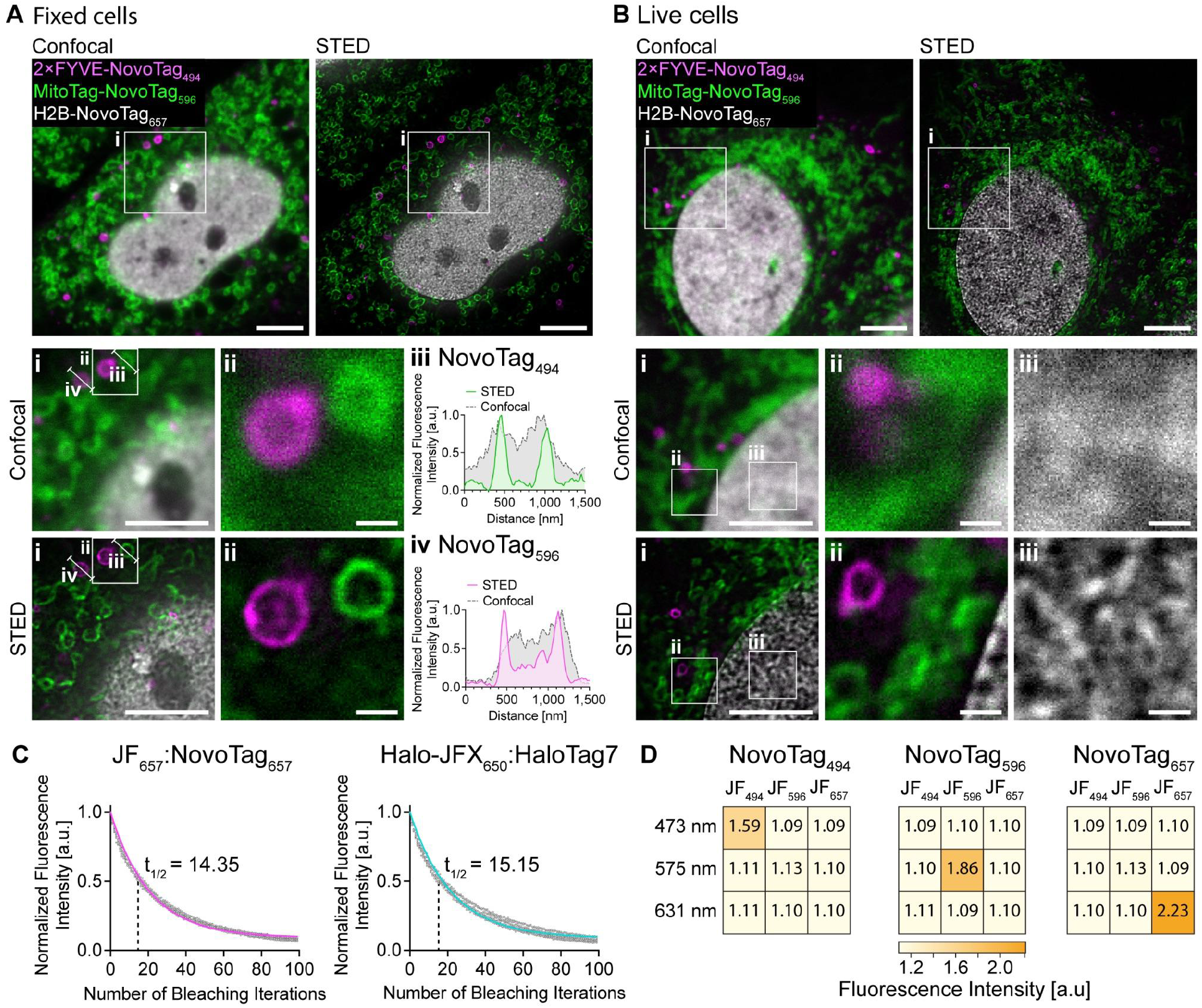
Multiplexed super-resolution fluorescence microscopy in live and fixed cells using NovoTags. **(A & B)** Multiplexed fluorescence microscopy in fixed (A) and live (B) HeLa cells: NovoTag_494_ labels endosomes (2×FYVE, magenta), NovoTag_596_ labels mitochondria (MitoTag, green), and NovoTag_657_ labels chromatin (H2B, white). Cells were stained with 50 nM of each dye (JF_494_, JF_596_, JF_657_) for 30 min at 37 °C, followed by three washing steps. Confocal and STED images (presented side by side at the top) were acquired on a Leica Stellaris 8 STED Falcon microscope. Bottom: numbered zoom-ins and line plots (ii, iv) for NovoTag_494_ (iii) and NovoTag_596_ (iv) for confocal and STED. **(C)** Photostability of JF dyes: HeLa cells expressing either NovoTag_657_ or HaloTag7 localized to mitochondria (MitoTag) were fluorescently labeled with 50 nM of each dye (JF_657_, Halo-JFX_650_) as above. Fluorescence intensity was measured after each bleaching iteration. For each sample, photobleaching of 10 individual cells was acquired in two independent experiments. All data points are shown (gray). Non-linear regression (one-phase decay) of the fluorescence signal was fitted. **(D)** Specificity of NovoTags: Each NovoTag (NovoTag_494,_ NovoTag_596,_ NovoTag_657_) was localized to mitochondria (MitoTag) in HeLa cells, labeled with one of each JF dye (JF_494,_ JF_596,_ JF_657_), and excited with one of three wavelengths (474 nm, 575 nm, 631 nm). Fluorescence intensity measurements were obtained from twelve fields of view for each condition in three independent experiments. For each combination, the mean fluorescence intensity is plotted. Scale bars: (A & B) 5 µm, zoom 1-2: 5 µm, zoom 3-6: 500 nm.

We evaluated the in cell specificity of the NovoTags by testing each combination of NovoTag (NovoTag_494_, NovoTag_596_, and NovoTag_657_), JF dye (JF_494_, JF_596_, and JF_657_), and excitation wavelength (473 nm, 575 nm, and 631 nm) in HeLa cells and quantifying the fluorescence intensities (**Fig. 2D, Fig. S11**). Only the specific combination of a matching NovoTag, JF dye, and excitation wavelength yielded a significant fluorescent signal, demonstrating high specificity in cells.

To explore the use of the NovoTags for multiplexed fluorescence imaging, we simultaneously targeted NovoTag_494_ to the early endosomal membrane (2×FYVE tag) (*18*), NovoTag_596_ to the inner nuclear membrane (emerin) (*19*, *20*), and NovoTag_657_ to the mitochondrial membrane (MitoTag) in HeLa cells. Cells were incubated with a dye cocktail consisting of 5 nM JF_494_, 20 nM JF_596_, and 5 nM JF_657_ for 15 min. After three washes, live imaging revealed clear signal separation (**Fig. S12**), demonstrating orthogonal and specific dye binding in cells and the multiplexing capability of this system.

Given the reported high photostability of JF dyes (*21*, *22*) and the observed high in-cell binding specificity of our binders, we evaluated the NovoTags for multiplexed super-resolution fluorescence microscopy using stimulated emission depletion (STED) imaging on fixed (**Fig. 2A, Fig. S13**) and live (**Fig. 2B**) HeLa cells. NovoTag_494_ was targeted to the early endosomal membrane (2×FYVE tag), NovoTag_596_ to the outer mitochondrial membrane (MitoTag), and NovoTag_657_ to chromatin (H2B) (*23*). Cells were incubated with a cocktail of all three JF dyes (50 nM each for 30 min). Following extensive washing, STED imaging revealed clear signal separation (**Fig. S13**) and enhanced resolution compared to confocal imaging (**Fig. 2A**). The combination of the three described NovoTags thus enables multiplexed cellular imaging in live, fixed, and super-resolution modalities.

### Tuning NovoTag_494_ for multiplexed fluorescence lifetime imaging microscopy (FLIM)

Our observation that the NovoTags modulate the fluorescence lifetimes of the JF dyes upon binding (**Fig. 1E**) suggests that the photophysical properties can be fine-tuned by modulating the designed binding pockets. As a proof of principle, we sought to design binders for JF_494_ with distinct, well-separated fluorescence lifetimes to enable multiplexed FLIM (*24*). We characterized dye binding and fluorescence intensity of 36 redesigns of the NovoTag_494_ binding pocket incorporating more hydrogen bonds, fewer π-π stacking interactions, and an altered electrostatic environment (**Fig. 3A**) and selected two designs NovoTag_494_^S^ (short lifetime) and NovoTag_494_^L^ (long lifetime) (**Fig. S14, Fig. 3B – D**). The excitation and emission profiles were similar to those of the original design; NovoTag_494_^L^ showed a slight blue-shift (**Fig. 3B – C**), NovoTag_494_^S^ exhibited slightly reduced brightness (**Fig. 3D**), and both displayed a modest decrease in binding affinity (**Fig. 3E**). Fluorescence lifetime measurements in HeLa cells showed a mean fluorescence lifetime of 2.00 ns (standard deviation (SD): 0.09 ns) for NovoTag_494_, 1.27 ns (SD: 0.25 ns) for NovoTag_494_^S^, and 2.78 ns (SD: 0.40 ns) for NovoTag_494_^L^ (**Fig. 3F**). These lifetime distributions were well separated, with Cohen’s *d* values of 4.5 (NovoTag_494_^S^:NovoTag_494_^L^), 3.9 (NovoTag_494_^S^:NovoTag_494_), and 2.7 (NovoTag_494_^L^:NovoTag_494_) (**Fig. 3F**).

**Fig. 3.**
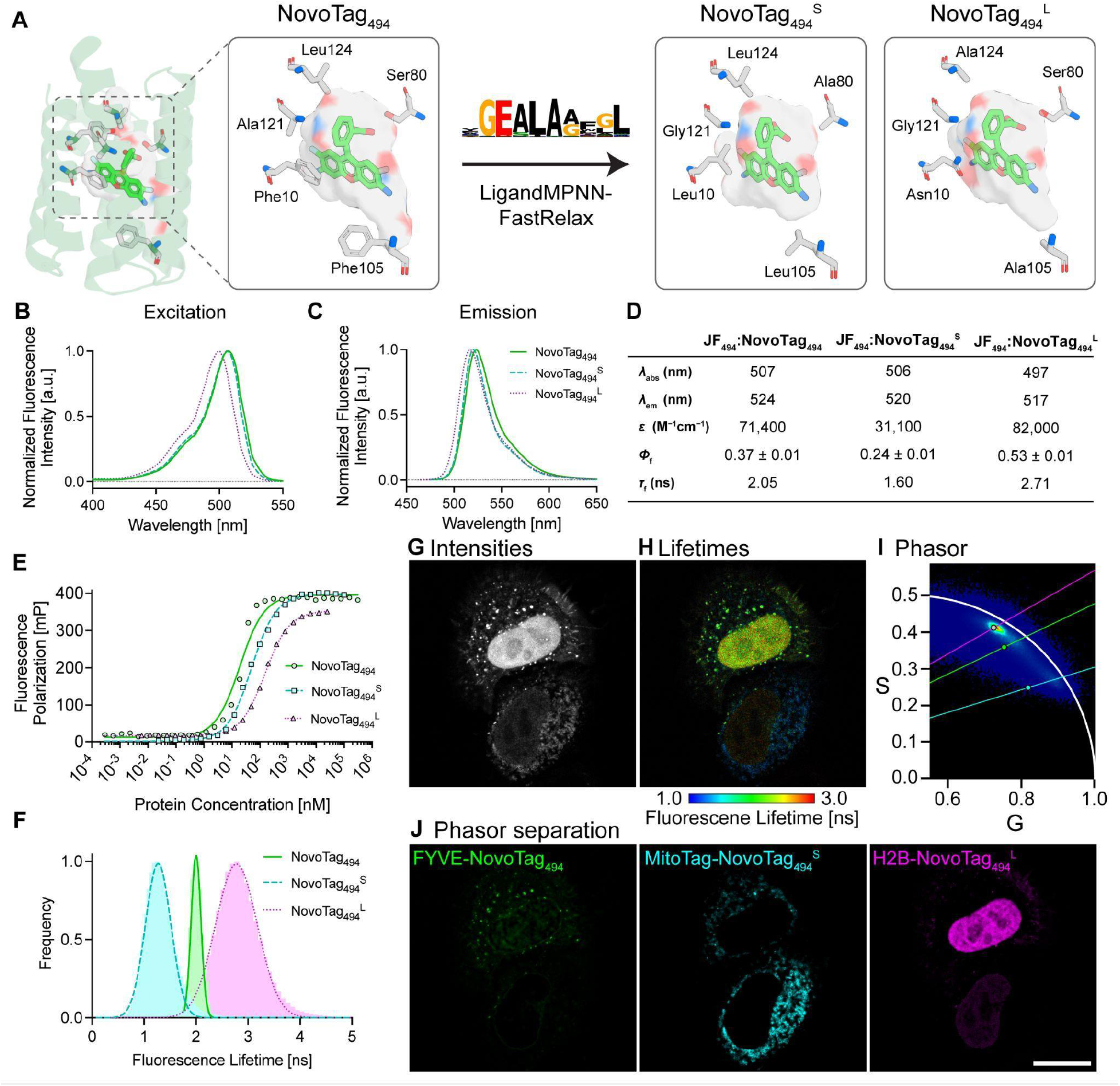
Fluorescence-lifetime-based multiplexed microscopy using NovoTags. **(A)** NovoTag_494_ binding site was subjected to sequence redesign using LigandMPNN-FastRelax, and two variants with different hydrogen-bonding networks, π–π-stacking profiles, and electrostatics of the binding pocket were selected with shorter or longer fluorescence lifetimes (NovoTag_494_^S^ and NovoTag_494_^L^). The sequence logo represents the relative frequency (letter height) of each amino acid in the redesigned residues. **(B & C)** Excitation (B) and emission (C) of JF_494_ dye bound to NovoTag_494_ and the two lifetime variants. **(D)** Photophysical characterization of the JF_494_-NovoTag_494_ variant conjugates. Parameters as specified in Fig. 1D. **(E)** Fluorescence polarization titration of NovoTag_494_, and of the two lifetime variants. A nonlinear regression (standard binding isotherm model) was fitted, and the K_d_ values were determined. NovoTag_494_^S^ and NovoTag_494_^L^ showed affinities to JF_494_ of 49.8 nM and 150.8 nM, respectively. For each sample, 3-4 independent measurements were acquired. **(F)** Distribution of fluorescence lifetimes of each NovoTag_494_ variant in cells. HeLa cells were transfected with one of the following plasmids: MitoTag-NovoTag_494_^S^, 2×FYVE-NovoTag_494_, or H2B-NovoTag_494_^L^, chemically fixed, and imaged in the presence of 10 nM JF_494_. For each sample, fluorescence images of 11-16 cellular regions were acquired. The histograms of fluorescence lifetimes are plotted, and fitted with a Gaussian distribution. **(G – J)** HeLa cells were co-transfected with MitoTag-NovoTag_494_^S^, 2×FYVE-NovoTag_494_, and H2B-NovoTag_494_^L^, chemically fixed, and imaged in the presence of 10 nM JF_494_ with a confocal microscope Leica Stellaris 8 Falcon. Shown are the total fluorescence intensity (G), the fluorescence lifetimes (H), and the phasor plot (I). Using phasor-based lifetime unmixing, the signals of the three labeled components were separated (J). Scale bar: 20 µm.

We evaluated multiplexed FLIM with these designs (**Fig. 3G – J**) by co-transfecting HeLa cells with MitoTag-NovoTag_494_^S^, 2×FYVE-NovoTag_494_, and H2B-NovoTag_494_^L^, followed by fixation and treatment with 10 nM JF_494_. Using phasor-based fluorescence lifetime unmixing, the signals of the three labeled components were separated. Signals showed specific signatures of mitochondria (MitoTag-NovoTag_494_^S^), endosomes (2×FYVE-NovoTag_494_), and chromatin (H2B-NovoTag_494_^L^), with minimal cross-talk between lifetimes, demonstrating successful fine-tuning of our NovoTags for multiplexed FLIM-based cellular imaging.

### Design of the covalent NovoTag_657_^cv^

Covalent dye binding is of particular interest for applications such as pulse-chase experiments and single-molecule tracking (*25–27*). We thus investigated the design of dye-specific covalent NovoTags. Fluorinated dyes such as JF_657_ can undergo nucleophilic aromatic substitution (S_N_Ar) on the fluorinated pendant phenyl ring (*28*). We surmised that placing a cysteine residue in NovoTag_657_ within close proximity to its electrophilic moiety would facilitate the formation of a covalent thioether bond between the dye and the protein (**Fig. S16A**).

We expressed and purified 47 such designs, incubated them with an excess of JF_657_ (10 µM for 12 h), and investigated the formation of a protein-dye covalent adduct using sodium dodecyl sulfate-polyacrylamide gel electrophoresis (SDS-PAGE). Twelve out of the 47 designs showed covalent binding (**Fig. S15, Fig. S16B**), and we named the design with the highest fluorescence intensity in the SDS-PAGE gel NovoTag_657_^cv^. Liquid chromatography-mass spectrometry (LC-MS; **Fig. S16C**) analysis revealed peaks corresponding to the labeled NovoTag_657_^cv^, confirming the formation of the covalent bond with the dye. LC-MS time-course analysis revealed 47.8% (SD = 1.6%) labeling after 3 h of incubation, suggesting a half-time (t_1/2_) of approximately 3 h for S_N_Ar conjugation kinetics (under excess JF_657_ conditions, **Fig. S16D**). Incubation of live *E. coli* or HeLa cells expressing NovoTag_657_^cv^ with JF_657_ for up to 18 h followed by SDS-PAGE analysis of whole-cell lysates (**Fig. S16E – F**) showed that the covalent bond forms in cells. Live fluorescence imaging of the HeLa cells confirmed colocalization between MitoTag-NovoTag_657_^cv^-eGFP and JF_657_ (**Fig. S17**), demonstrating the feasibility of covalent labeling with the NovoTags for cellular imaging.

### NovoSplit enables chemically induced dimerization

Chemically induced dimerization (CID) systems have been widely used to promote protein–protein interactions for applications ranging from signal transduction and gene expression to protein localization and degradation (*29–32*). We reasoned that designed split variants of the NovoTags could function as a CID system, wherein the fluorescent JF dye functions to induce dimerization within live cells, allowing for precise control of protein interactions and downstream processes, while simultaneously providing a direct fluorescent readout of the induced interactions.

We designed split versions of NovoTag_657_, and 47 designs predicted to assemble in the presence of dye were evaluated in a mammalian two-hybrid screen for minimal self-dimerization in the absence of the dye and inducible dimerization upon addition of JF_657_ (**Fig. S18**). The five designs exhibiting the highest dye-induced reporter activation were further evaluated by fluorescence colocalization analysis in HeLa cells (**Fig. S19**; the best of these split designs (**Fig. S19B**) was derived from the top non-covalent design NovoTag_657_ (**Fig. 1C**)). To reduce residual self-dimerization in the absence of JF_657_ (**Fig. S19B**), we eliminated a salt bridge between Glu14 and Arg53 (**Fig. S20A**) by introducing the Arg53Ala substitution; this eliminated the non-induced dimerization, while induction of dimerization by JF_657_ was not impaired (**Fig. S20B & C**). We named this split system NovoSplit_657_. The AF3 prediction of NovoSplit_657_ aligned closely with the holo crystal structure of NovoTag_657_ (Cα-RMSD = 0.38 Å) (**Fig. 4B**).

**Fig. 4.**
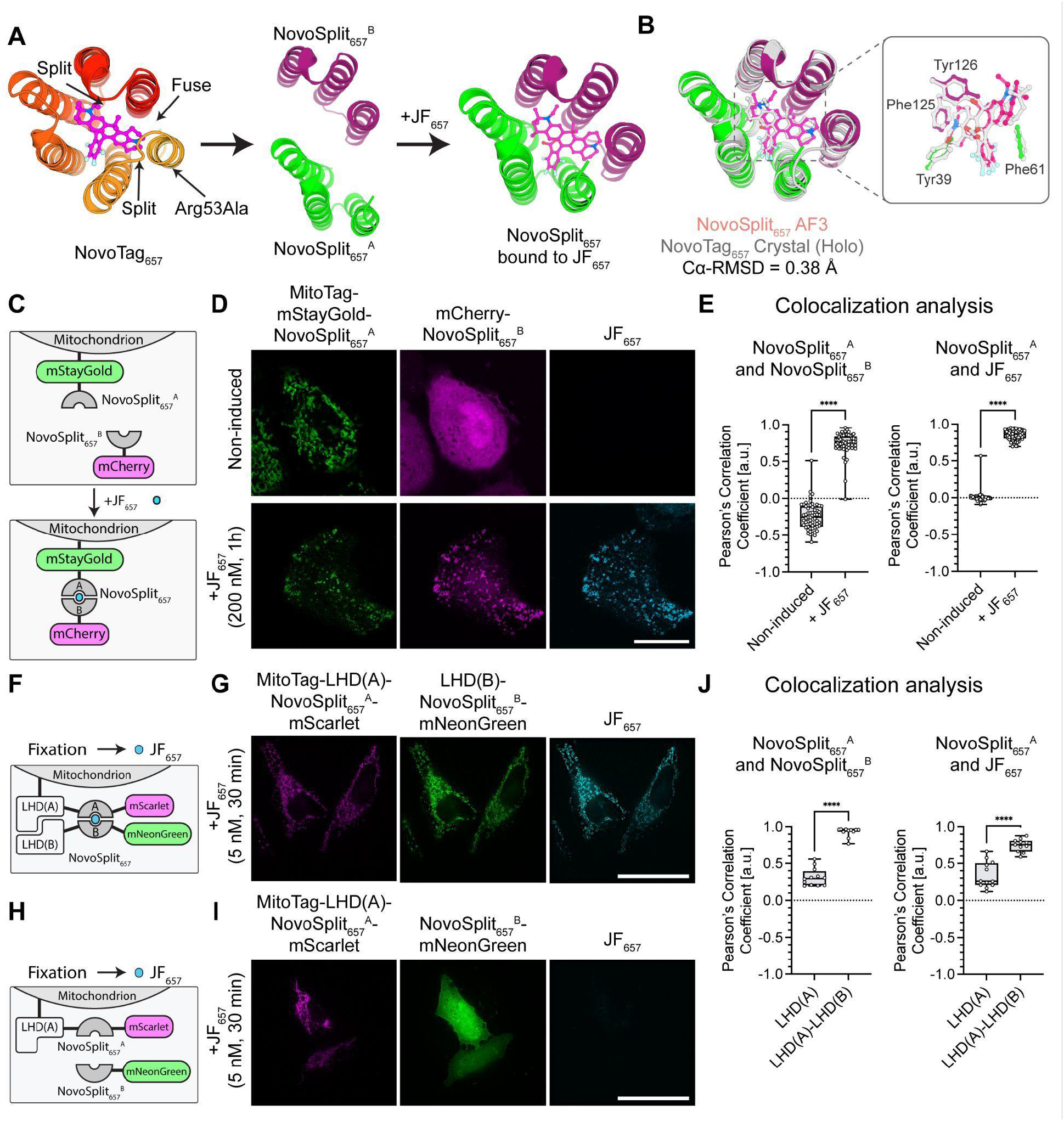
Chemically induced dimerization and proximity biosensing using NovoSplit_657_. **(A)** The design of NovoSplit_657_ is based on NovoTag_657_ that was split at two sites, and the original C- and N-termini were fused with a short linker. The fluorescent dye JF_657_ induces dimerization of the two monomers NovoSplit_657_^A^ and NovoSplit_657_^B^. To prevent non-induced dimerization, an Arg53Ala substitution was introduced into NovoSplit_657_^B^. **(B)** The NovoSplit_657_ AF3 prediction (in color) superimposed with the crystal structure of NovoTag_657_ (gray). **(C – E)** NovoSplit_657_ as a CID system: HeLa cells were transfected with MitoTag-mStayGold-NovoSplit_657_^A^ and mCherry-NovoSplit_657_^B^ (C). Cells were incubated with 200 nM JF_657_ for 1 h, washed three times, chemically fixed, and imaged with a confocal microscope Zeiss LSM 980 AiryScan (D). Colocalization was analyzed by calculating the Pearson’s correlation coefficient between NovoSplit_657_^A^ and NovoSplit_657_^B^, as well as between NovoSplit_657_^A^ and JF_657_ (E). For each sample, 59 (non-induced) and 61 (+JF_657_) individual cells were analyzed. **(F – J)** NovoSplit_657_ as proximity biosensor: HeLa cells were transfected with MitoTag-LHD(A)-NovoSplit_657_^A^-mScarlet and either LHD(B)-NovoSplit_657_^B^-mNeonGreen (F) or NovoSplit_657_^B^-mNeonGreen (negative control, H). Cells were chemically fixed and fluorescently labeled with 5 nM JF_657_ for 30 min, followed by three washing steps. Fluorescence images were acquired on an OMX SR microscope (G & I). For experiment in G, colocalization was analyzed by calculating the Pearson’s correlation coefficient between NovoSplit_657_^A^ and NovoSplit_657_^B^, as well as between NovoSplit_657_^A^ and JF_657_ (J). For each sample, 11 (LHD(A)-LHD(B)) and 12 (LHD(A)) individual cells were analyzed. All data are shown as Box and Whiskers plots indicating the median (center of box), 25% and 75% quartiles (bounds of box), minimum and maximum values (bars). Each data point represents one cell. Unpaired two-sided Welch’s t-tests were conducted. Statistical significance: ∗∗∗∗ p < 0.0001. Scale bars: (D) 20 µm, (G & I) 40 µm.

We evaluated NovoSplit_657_ as a CID system in HeLa cells. One split fragment, NovoSplit_657_^A^, was targeted to the outer mitochondrial membrane using a MitoTag and additionally fluorescently labeled with mStayGold (*33*). The second split fragment, NovoSplit_657_^B^, was fluorescently labeled with mCherry (*34*) and expressed as a cytosolic protein (**Fig. 4C**). Incubation with JF_657_ (200 nM for 1 h) induced translocation of mCherry-NovoSplit_657_^B^ from the cytosol to the mitochondria, with a Pearson’s correlation coefficient (r) between mStayGold and mCherry of 0.74 (SD: 0.15). The JF_657_ fluorescent signal colocalized with both mStayGold-NovoSplit_657_^A^ and mCherry-NovoSplit_657_^B^ (r = 0.78, SD: 0.06, and r = 0.88, SD: 0.05), demonstrating that the small molecule both induces dimerization and provides a fluorescent readout of the newly-formed interaction (**Fig. 4D & E**). Titration with JF_657_ showed that concentrations as low as 50 nM induce dimerization, with higher dye concentrations yielding faster kinetics (**Fig. S21**). Live-cell fluorescence imaging with 1 µM JF_657_ showed a half-time (t_1/2_) of 5.8 min (95% confidence interval (CI): 5.0-6.7 min) (**Fig. S22, Movie S2**). Evaluation of the dye binding specificity of NovoSplit_657_ showed that only JF_657_ induced dimerization, whereas JF_494_ and JF_596_ did not (**Fig. S23**).

### NovoSplit enables native protein-protein proximity biosensing

We hypothesized that NovoSplit_657_ could be additionally used to probe native protein-protein interactions, similar to bimolecular fluorescence complementation systems (*35*) such as split-GFP, if the two components of NovoSplit_657_ were fused to two proteins of interest, and the JF_657_ dye was only introduced after cell fixation. We reasoned that if the proteins of interest were in close proximity within a cell, small rearrangements after fixation could occur to enable JF_657_ dye-facilitated dimerization of the split system, thereby providing a direct fluorescent readout of protein proximity. If, on the other hand, the two proteins of interest were far apart, fixation would prevent the large scale rearrangements required for assembly of the two halves of the split system, and no fluorescence would be observed.

To evaluate protein-protein proximity biosensing using the NovoSplit system, we used a *de novo*-designed protein pair, LHD(A) and LHD(B), which forms stable heterodimers (*36*). LHD(A) was fused to NovoSplit_657_^A^-mScarlet and targeted to the outer mitochondrial membrane using a MitoTag. LHD(B) was fused to NovoSplit_657_^B^-mNeonGreen and expressed as a cytosolic protein. As LHD(A) and LHD(B) spontaneously form heterodimers, we expected that both constructs would colocalize, bringing NovoSplit_657_^A^ and NovoSplit_657_^B^ in proximity (**Fig. 4F**). Following chemical fixation and incubation with JF_657_ (5 nM for 30 min), fluorescence microscopy showed colocalization of mScarlet and mNeonGreen (r = 0.93, SD: 0.06), indicating formation of the LHD(A)-LHD(B) heterodimer, and a clear JF_657_ fluorescence signal that colocalized with both fluorescent proteins (r = 0.74, SD: 0.09; r = 0.79, SD: 0.08) (**Fig. 4G & J**), showing that the NovoSplit_657_^A^ -NovoSplit_657_^B^ -JF_657_ ternary complex can assemble after fixation. As a control, we repeated the same experiment, expressing NovoSplit_657_^B^ fused to mNeonGreen but omitting LHD(B) (**Fig. 4H**). Without the LHD(A)-LHD(B) pair to drive association, the NovoSplit_657_^B^-mNeonGreen was distributed throughout the cytosol, and the two constructs showed minimal colocalization (r = 0.31, SD: 0.12), and no localized JF_657_ fluorescence (**Fig. 4I** & **J**). These data confirm our conjecture that addition of dye post-fixation only leads to reconstitution of the fluorescent ternary complex if the two halves of the split system are already in close proximity. Thus, by adding dye post-fixation, the NovoSplit system can be used to detect interactions between two proteins of interest without introducing spurious associations; the fluorescent signal provides a snapshot of the extent of association at the time of fixation, rather than the cumulative amount of complex formed as with some irreversible split fluorescent protein systems (*37*).

## Discussion

Our NovoTags combine the superior photophysical properties of small-molecule fluorophores with the genetic programmability of proteins. Because the designed binding sites are dye-specific, NovoTags enable the direct and simultaneous use of multiple bright, photostable, non-modified JF dyes in the same cell for multiplex imaging; in contrast, the shared conjugation chemistry of HaloTag (*5*) or SNAP-tag (*6*) only allows use of one dye with each tag in a single experiment. In addition, the smaller size of NovoTags should reduce steric hindrance and potential perturbation to the tagged proteins. The tunability of the NovoTag binding sites further expands opportunities for advanced microscopy and approaches to modulate or measure protein interactions in cells: the covalent NovoTag_657_^cv^ offers advantages in applications requiring permanent labeling, including pulse-chase experiments, single-molecule (*25–27*), and *in vivo* imaging. NovoTags that modulate fluorescent lifetimes extend imaging beyond spectral unmixing, enabling multiplexed fluorescence lifetime imaging microscopy (FLIM) that is especially useful for live-cell imaging, as reducing the number of spectrally distinct channels reduces photobleaching. NovoSplit converts dye binding into conditional protein dimerization with high specificity and a direct fluorescent readout. With the fluorophore introduced after fixation, NovoSplit functions as a very sensitive (due to the brightness of the JF dyes) and low background native interaction proximity sensor.

The design strategy described here should be extendable to a wider range of synthetic fluorophores, including near-infrared dyes and fluorophores with specialized properties such as photoactivation or blinking (*21*, *38*, *39*). Given that advanced spectral unmixing can achieve spectral resolution for fluorescent dyes with emission maxima separated by 25 nm (*40*, *41*), it should be possible to develop 10 or more resolvable NovoTag–dye combinations spanning the entire spectrum. The additional capability of separation along the orthogonal lifetime dimension, demonstrated here with the three lifetime-resolvable NovoTag_494_ variants, could expand such a panel to 3 × 10 = 30 or more simultaneously resolvable probes; being able to thus simultaneously resolve and track many different tagged proteins in the same cell or tissue could enable considerable advances in molecular cell biology. Together with recent related studies (*42–44*), our work outlines a general framework for integrating synthetic chemistry with protein engineering to advance biological imaging and related fields.

## Acknowledgement

Crystallographic diffraction data were collected at the Advanced Photon Source (APS) and the National Synchrotron Light Source II (NSLS-II). At the APS, this work used the Northeastern Collaborative Access Team (NE-CAT) beamlines, funded by the National Institute of General Medical Sciences (NIGMS) from the National Institutes of Health (NIH) (P30 GM124165), utilizing the Eiger 16M detector on beamline 24-ID-E (NIH-ORIP HEI grant S10OD021527). This research was performed on APS beam time award (https://doi.org/10.46936/APS-192610/60015997) from the APS, a U.S. Department of Energy (DOE) Office of Science User Facility operated by Argonne National Laboratory under Contract No. DE-AC02-06CH11357. At NSLS-II, this work used beamline 17-ID-2 (FMX), supported by the Center for BioMolecular Structure (CBMS). CBMS is primarily supported by the NIH/NIGMS through a Center Core P30 Grant (P30GM133893) and by the DOE Office of Biological and Environmental Research (KP1605010). NSLS-II is a U.S. DOE Office of Science User Facility operated for the DOE Office of Science by Brookhaven National Laboratory under Contract No. DE-SC0012704. This work was delivered in part as part of the MATCHMAKERS team supported by the Cancer Grand Challenges partnership funded by Cancer Research UK (CGCATF-2023/100008), the National Cancer Institute (OT2CA297288), and the Mark Foundation for Cancer Research. We thank the EMBL Advanced Light Microscopy Facility, especially Martin Fritsch, for support with fluorescence microscopy. We thank David Agard, Helen Eisenach, Shingo Honda, Thomas Schlichthaerle, Avi Swartz, Jonas Wilhelm, Xinru Wang, Yulai Liu, Hojun Choi, Green Ahn, Mohamad Abedi, Stephanie Hanna, Nancy Zhu, Todd J. Martinez, and Ariana Tkachuk for helpful discussions and comments during the preparation of this manuscript.

## Funding

The research is supported by the Howard Hughes Medical Institute (GR020267: C19 HHMI INITIATIVE - 66-0656 - 2021, to A.K., E.J., N.R.; Janelia Research Campus to T.A.B, R.P., J.B.G., L.D.L.); Merck (GR024322: MERCK COLLABORATION - 66-8270 - 2021, to L.T., D.V.); The Audacious Project at the Institute for Protein Design (PG117878: Audacious Hub, PG117866: Audacious Discretionary Sub, to D.B.); European Commission through the ARISE programme (Horizon 2020 Research and Innovation Programme under the Marie Skłodowska-Curie grant agreement number 945405, to S.K.); Erwin Schrödinger Postdoctoral Fellowship (J-4663, to E.M.); Chan Zuckerberg Initiative (Grant Number 2023-332009, to J.M and D.B.); and the Damon Runyon Cancer Research Foundation (DRG-2507-23 to W.C.).

## Author contribution

Long Tran: project design and execution, manuscript preparation; Steffen Klein: project design, pipeline design for split binder, cellular super-resolution imaging, figure and manuscript preparation; Shajesh Sharma: library preparation; David Juergens: computational scripts collaboration; Justin Decarreau: cellular imaging, analysis, and discussion, manuscript preparation; Bingxu Liu: discussion; Yujia Wang: pipeline design; Asim Bera: solved crystal structure; Xinting Li: LC-MS data curation; Alex Kang: xtal curation; Dionne K Vafeados: library preparation; Nicole Roullier: library preparation; Yang Bo: experimental preparation; Edin Muratspahić: experimental preparation; Wei Chen: experimental preparation; Gyu Rie Lee: design pipeline and experimental preparation; Timothy A. Brown: stop-flow experiments; Ronak Patel: fluorescence lifetime measurements; Jonathan B. Grimm: chemical synthesis; Luke D. Lavis: spectroscopy and project design; Julia Mahamid: project design, imaging supervision, and manuscript preparation; Linna An: project design, computational analysis, and manuscript preparation; David Baker: project design and manuscript preparation.

L.T. and S.K. contributed equally as first authors. D.J., S.S., and J.D. contributed equally as second authors, and everyone agrees that their authorship order can be exchanged to benefit their own career development.

## Conflict of Interest

US patent 12,344,594 and US patent application US20260001857 describing fluorophores and variant compositions (with inventors J.B.G. and L.D.L.) are assigned to HHMI. L.D.L. is a scientific co-founder and shareholder of Eikon Therapeutics. L.T., S.K., L.A., J.D., L.D.L, J.M., and D.B. have filed a provisional patent application 64/041,005 that incorporates discoveries described in this article. All authors declare no conflict of interest.

## Data, code, and materials availability

Crystal structures of NovoTag_657_ are available at the RCSB Protein Data Bank (PDB) with accession numbers 9PVL for the apo structure and 11MQ for the holo structure. Precommercial samples of JF_494_, JF_596_, and JF_657_ can be requested at https://dyes.janelia.org. Plasmids generated in this work (listed in Table S3) are available at Addgene with accession numbers 255873, 255874, 255875, 255876, 255877, 255878, 255879, 255880, 255881, 255882, 255883, 255884, and 255885. The pseudocycle Cα RFdiffusion code is available on GitHub (https://github.com/baker-laboratory/CA_RFDiffusion); the design pipeline is available on GitHub (https://github.com/steffen-klein/novotag-design) and archived on Zenodo (*45*). Fluorescence microscopy and SDS-PAGE raw data are archived on Zenodo (*46*).

## Supplementary Materials

- Materials and methods
- References
- Figures S1 – S23
- Tables S1 – S3
- Movies S1 – S3
- Reproducibility Checklist

## Supplementary materials

## Material and methods

### Computational design pipeline for dye binder using pseudocycle Cα RFdiffusion

Ligand rotamers were generated using CREST (*47*). We developed a ligand-aware symmetry-conditioned implementation of Cα (C-alpha of protein backbone) RFdiffusion (*10*), namely pseudocycle Cα RFdiffusion, to generate protein backbones around defined ligands. Given a choice of symmetry (N) and symmetry unit length (L, the full protein length will be LxN), at each denoising step (*i.e.,* protein generation step), we symmetrize the RoseTTAfold pair representation by collecting all L by L length blocks of features corresponding to a unique subunit-subunit interaction, aggregating them into a single block. To enforce structural symmetry in the pair representation, the full L×L interaction matrix is partitioned into N×N sub-blocks according to the user-defined repeat unit length, and a symmetry class label is assigned to each RoseTTAfold block pair. Blocks sharing the same class label, which correspond to geometrically equivalent inter- and intra-repeat interactions, are aggregated by computing their element-wise mean, which is then updated back to all equivalent block positions. This operation imposes repeat symmetry prior to the subsequent structural update step. Analogously, symmetry can be enforced by replacing each group of class-equivalent sub-blocks with their element-wise maximum, rather than their mean. Element-wise maximum operation preferentially retains the most strongly activated features across all symmetry-related subunits (according to RoseTTAfold pair representation output). This operation allows the block in which a particular pairwise interaction signal is most pronounced to provide the most informative representation to be propagated.

Designed backbones, from pseudocycle Cα RFdiffusion, were then filtered by ligand surface area, retaining only those with a satisfactory contact area. Sequences were designed using LigandMPNN-FastRelax (*9*). Protein folding and ligand binding were evaluated using Rosetta (*13*) and AF2 (*14*), and the results were filtered using previously published selection criteria (*9*). Overall, the major filters used are: pLDDT >= 0.85, Cα-RMSD <= 1.5, and pTM >= 0.6. We further selected designs that ranked in the 70th percentile or higher for contact molecular surface area, and in the lowest 30th percentile for ddG.

For further details on the pseudocycle Cɑ RFdiffusion design pipeline, see the code repository on GitHub (https://github.com/steffen-klein/novotag-design) or Zenodo (*45*).

### Yeast surface display library preparation

All experiments were performed following previously published protocols (*9*). All library oligos were obtained from Twist Bioscience. Gene fragments were sourced from Twist Bioscience or IDT. The following materials were used: KAPA HiFi HotStart Uracil+ReadyMix (Roche, #KK2802), KAPA HiFi HotStart Kit (Roche, #KK2101), EvaGreen Dye (Biotium, #31000), USER enzyme (NEB, #M5508), NEBNext End Repair Module (NEB, #E6050L). Chicken Anti-C-Myc antibody - FITC conjugated (ICL, #CMYC-45F), mouse anti-C-Myc antibody - PE conjugated (Cell Signaling Technology, 9B11, #3739), mouse anti-C-Myc antibody - Alexa Fluor 647 conjugated (BioLegend, E10, #626810). DNA purification kits were purchased from QIAgen. Unless otherwise specified, all chemicals and consumables were purchased from Thermo Fisher Scientific.

Designed protein sequences were back-translated with DNAworks (*48*), ensuring that common restriction enzyme cut sites were excluded. Each nucleotide sequence was split into two segments, a 5′ segment and a 3′ segment, with complementary overlap in the center; this was automated using an in-house Python script. To minimize qPCR bias from oligo-length variation, designs were binned into groups of ∼1,000 by length. Primers matching the pETCON3 yeast-surface-display vector were appended to the 5′ end of the 5′ segment and the 3′ end of the 3′ segment, while inner primers containing AA or TT were added to the opposite ends to streamline library PCR. Depending on the design length, 250 or 300 nt oligo pools were ordered from Twist Bioscience.

Oligo pools were resuspended in water to 100 ng/µl, stored at −20 °C, and then diluted to 2.5 ng/µl for qPCR. Individual 5′ and 3′ segments were amplified with custom primers using KAPA HiFi HotStart Uracil+ReadyMix, with DNA purification between qPCR steps. USER enzyme and the NEBNext End Repair Module removed inner primers per the manufacturer’s protocols. After purification, the digested 5′ and 3′ fragments were assembled by the same qPCR protocol, and assemblies were verified by gel electrophoresis. Desired DNA bands were excised and purified for further amplification. Approximately 4-6 µg of the pooled oligos were used to transform electrocompetent yeast. The oligo pool and linearized pETCON3 vector were mixed at a 3:1 ratio, and yeast were transformed by electro-transfer following the previously published protocol (*9*).

### Yeast surface display analysis for enrichment of potential binders

The established yeast-surface-display protocol (*9*) was followed with only minor adjustments. PBSF (PBS containing 0.1 % w/v BSA) served as the buffer for every yeast-sorting step. Strain EBY100 was used throughout all surface-display experiments. Library fluorescence-activated cell sorting (FACS) was conducted on a Sony SH800S cell sorter fitted with a 100 µm chip and 488 nm / 561 nm / 638 nm lasers.

Each yeast library was first subjected to an expression sort to enrich for binders with adequate surface display. Binder-expressing cells were stained 1:50 (v/v) with anti-cMyc-PE for 30 min at 4 °C; cells exhibiting elevated PE signal were collected for downstream binding assays. These cells were regrown in C-Trp-Ura medium with 2% glucose (CTUG) and then induced overnight at 30 °C in SGCAA medium.

The induced yeast was stained 1:35 (v/v) with anti-cMyc-PE for 45 min at 4 °C, washed twice in PBSF, and incubated with the dye ligand for 60-120 min at 4 °C. After two further PBSF washes, cells were sorted. Unstained induced yeast served as a negative control; cells positive in both dye fluorescent channel and PE channel above this control were collected and expanded in CTUG. After 1-2 days of growth, cultures were re-induced in SGCAA medium and sorted again as described. Typically, two to three rounds were required to achieve clear enrichment, and when binding populations exceeded 2%, titrations were performed to identify the strongest binders.

Samples from the expression step and every sorting round were reserved for MiSeq or NextSeq 2000 analysis. Yeast-sorting data were processed and visualized using FlowJo v10 (FlowJo Inc.).

### Expression and purification of binders

The workflow largely followed an earlier publication (*9*), with all reagents and consumables obtained from Thermo Fisher Scientific unless noted otherwise. Selected designs were back-translated with DNAWorks, and eblocks (IDT or Twist Bioscience) were inserted into a pET29b-based vector bearing a C-terminal, SNAC-cleavable His-tag by Golden Gate assembly (New England Biolabs) before transformation into BL21 competent *E. coli*.

### Small-scale expression and purification of verified binders

Designs exhibiting promising activity were subjected to rapid, small-scale expression and purification following previous protocols (*49*, *50*). 1 µl Golden Gate subcloning reactions were assembled in 96-well PCR plates, transformed into chemically competent BL21(DE3) cells, and, after a 1 h outgrowth, distributed into four 96-deep-well plates, each containing 1 ml autoclaved auto-induction medium (Terrific Broth supplemented with 0.05 mg/ml Kanamycin, 2 mM MgSO₄, and 1 × 5052 (0.5% glycerol, 0.05% glucose, and 0.02% α-lactose monohydrate)), for a final culture volume of ∼4 ml per construct. After 20-24 h of incubation, cells were harvested and lysed, and the clarified lysate was loaded directly onto 100 µl Ni-NTA agarose beds in 96-well fritted plates pre-equilibrated with Tris wash buffer. Following thorough washing, bound proteins were eluted in 200 µl Tris buffer containing 500 mM imidazole.

Eluates were sterilized through a 0.22 µm 96-well filter plate (Agilent, #203940-100) before size-exclusion chromatography (SEC). Samples were analyzed on an ÄKTA FPLC equipped with an autosampler compatible with 96-well source plates. Monomeric binders were resolved on a Superdex 75 Increase 5/150 GL column (Cytiva, #29148722).

### Large-scale expression and purification of verified binders

For designs with validated binding and catalyzing activities, we perform large-scale expression and purification for detailed characterization.

Cultures (50 ml) were grown in auto-induction Terrific Broth within 250 ml baffled Erlenmeyer flasks for 6 h at 37 °C, followed by 24 h at 18 °C, shaken at ∼200 × g in New Brunswick Innova 44 shakers. Cells were harvested, resuspended in 25 ml lysis buffer (25 mM Tris, 100 mM NaCl, pH 8, a protease-inhibitor tablet), and sonicated for 4.5 min (10 s on / 10 s off, 65% amplitude). After a 30 min spin at 14,000 × g, the supernatant was incubated for 1 h with 1 ml Ni-NTA resin (Qiagen) in a BIO-RAD Econo-Pac gravity column, rotating gently. Resin washes comprised 20 ml of low-salt buffer (50 mM Tris, 100 mM NaCl, 10 mM imidazole, pH 8) and 20 ml of high-salt buffer (50 mM Tris, 1,000 mM NaCl, 10 mM imidazole, pH 8). Proteins were eluted with 2 ml of elution buffer (20 mM Tris, 100 mM NaCl, 500 mM imidazole, pH 8) and polished on a Superdex 75 Increase 10/300 GL column (ÄKTA system) in isocratic TBS (20 mM Tris, 100 mM NaCl, pH 8).

For proteins used in fluorescence polarization and crystallography experiments, His-tags were removed via on-bead SNAC cleavage. After binding, columns were rinsed twice with lysis buffer to eliminate imidazole, equilibrated with 10 ml SNAC buffer (100 mM CHES, 100 mM acetone oxime, 100 mM NaCl, 500 mM guanidine chloride, pH 8.5), and then incubated overnight at room temperature with 20 ml fresh cleavage solution (2 mM NiCl₂ in SNAC buffer). Resin was washed with 10 ml weak-elution buffer (20 mM Tris, 100 mM NaCl, 10 mM imidazole, pH 8); both the cleavage and wash fractions were collected and assessed by SDS-PAGE for tag-free protein. These proteins were pooled, concentrated, subjected to the same SEC protocol as above, and verified by mass spectrometry.

### Fluorescence polarization (FP) assay for dye binder affinity of binders

The protein concentrations were measured using UV-Vis. All FP studies and analyses were performed following a previously established protocol (*9*) on a Biotek Synergy Neo2 Reader equipped with a Dual PMT optical filter cube (#1035108, #1035065, #1035064). The proteins were serially diluted, and 0.8 nM of dye ligands were added to the diluted protein in 20 mM Tris, 100 mM NaCl, pH 8 buffer. The mixtures were transferred to a microplate, Corning 96 well half area flat bottom non-binding surface black plate (Corning, #3686). Samples were serially diluted within the microplate and allowed to rest at ambient temperature for 5 min. The sample plates were then incubated on a vibrating shaker (Heidolph Titramax 1000 Vibrating Shaker) at 1,000 rpm, 25 °C, for an additional 25 min prior to FP reading. The parallel and perpendicular light were read with scale to the highest-intensity well. At least three independent FP were performed as replicates. FP signal was calculated based on the following equation:

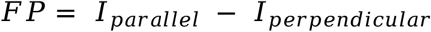

The data were fitted using a standard binding isotherm model with baseline offset using GraphPad Prism described by:

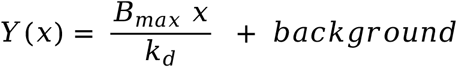

### Stopped-flow assay for NovoTag_494_ binding kinetics analysis

To measure dye-protein binding rates, we used a stopped-flow procedure where equimolar amounts of purified dye binding protein and JF_494_ dye were mixed, with association monitored by fluorescence polarization. Protein and dye were mixed in 20 mM Tris, 100 mM NaCl, pH 7.5 to yield varied amounts of dye:protein mixes. Stopped flow experiments were performed at 37 °C using a model SX20 Applied Photophysics stopped-flow spectrometer in fluorescence polarization configuration with excitation from a 490 nm LED with fluorescence emission detection through 515 nm long pass filters for both parallel and perpendicular polarized light signals. Various concentrations of binding solution mixes were chosen based on the K_d_ values from *in vitro* binding curves. Three replicates were performed for each concentration. Kinetics were analyzed using a full reversible bimolecular model. Under equimolar conditions, the complex formation over time is described by:

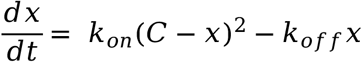

The anisotropy varies linearly with the fraction of dye bound to the protein:

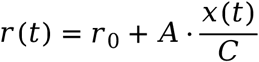

Where *r(t)* is the observed anisotropy, *r_o_* is the baseline anisotropy, and *A* is the amplitude.

All traces were fitted using a nonlinear least squares optimization, and global fitting was done with optimization using trust-region reflective algorithms.

### *In vitro* photophysical characterization of JF dyes and JF dye complexes/conjugates

The spectral properties of free dyes and NovoTag complexes were measured in 20 mM Tris, 100 mM NaCl buffer, pH 8 (TBS). The spectral properties of JF_657_-HaloTag ligand bound to HaloTag7 protein were measured in 10 mM HEPES, pH 7.3, containing 0.1 mg/ml CHAPS. All measurements were taken at ambient temperature (22 ± 2 °C) using 1 cm path length, 3.5 ml quartz cuvettes (Starna Cells). Dyes were prepared as stock solutions in DMSO and diluted such that the DMSO concentration was less than 0.1% (v/v). NovoTag proteins were added in excess to ensure >97% binding. HaloTag7 protein was added in 2-fold excess and incubated until a consistent absorption spectrum was observed (>60 min). Maximum absorption wavelength (*λ*_abs_) and extinction coefficients (*ε*) of dyes and their complexes/conjugates were measured on a Cary Model 4000 spectrometer (Agilent). Absorption spectra are averages (*n* = 3), and normalized spectra are shown for clarity. Fluorescence spectra and maximum emission wavelength (*λ*_em_) of the dye and their complexes/conjugates were measured on a Cary Eclipse fluorometer (Agilent); normalized spectra are shown for clarity. Absolute fluorescence quantum yield values (*Φ*_f_) of the dyes and their complexes/conjugates were measured using a Quantaurus-QY spectrometer (C11374, Hamamatsu). Measurements were carried out using dilute samples (*A* < 0.1), and self-absorption corrections were performed using the instrument software. Reported *Φ*_f_ values are averages of measurements from two separate dye solutions.

### *In vitro* fluorescence lifetime measurements

For fluorescence lifetime measurements, purified protein (88 μM) was mixed with dye (2 μM) in HEPES buffer (10mM HEPES, 0.1 mg/ml CHAPS, and 100 mM NaCl at pH 7.3) and incubated at RT for 8 h. A 100 μl aliquot of the protein–dye mixture was transferred to a CellVis 96-well glass-bottom plate for imaging. FLIM measurements were performed on a SP8 FALCON microscope (Leica Microsystems) equipped with a tunable pulsed white light laser (470–670 nm) operating at 20 MHz. Images were acquired using a 40×/1.30 NA oil-immersion objective (HC PL APO, Leica Microsystems) and a HyD SMD 2 detector (Leica Microsystems). Adaptive focus control was used during image acquisition. Fluorescence lifetime data were analyzed using LAS X FLIM software (Leica Microsystems). Lifetime decays were fitted using either a one-component or two-component exponential model as appropriate. The amplitude-weighted average fluorescence lifetime is reported for three independent measurements.

### Circular dichroism spectra analysis

Circular dichroism (CD) spectra for the chosen designs were acquired using a Jasco J-1500 spectrometer. Each sample (0.5 mg/ml) was prepared in TBS at pH 8 and placed in a 1 mm path length cuvette. Raw CD data were converted to molar mean residue ellipticity by dividing the signal by N × C × L × 10, where N is the residue count, C is the protein concentration, and L is the optical path length (0.1 cm).

### Crystallography of NovoTag_657_-apo and NovoTag_657_-holo

Crystallization experiments were conducted using the sitting drop vapor diffusion method. Initial crystallization trials were performed via sitting-drop vapor diffusion in 96-well plates at 20 °C, using a mosquito LCP liquid handler (SPT Labtech) to dispense 200 nl drops. Plates were imaged using UVEX microscopes and UVEX PS-256 from JAN Scientific. For NovoTag_657_-apo, diffraction quality crystals formed in 0.2 M ammonium sulfate, 0.1 M Tris pH 7.5, and 20 % (w/v) PEG 5000 MME in 11 % (w/v) PEG 3350, 0.1 M HEPES pH 7.5. For NovoTag_657_-holo, crystals formed in 0.2 M Lithium sulfate, 0.5 M Zinc acetate dihydrate, 0.1 M BisTris pH 7.5, 22.5% (v/v) PEG Smear Broad (PEG Smear Broad contains: PEG1000, PEG10000, PEG2000, PEG3350, PEG400, PEG4000, PEG600, PEG6000, PEG8000, PEG methyl ether 500 and PEG methyl ether 5000). We added 0.5 µl of 2 mM JF_657_ dye to the drops containing NovoTag_657_-apo crystals and allowed them to soak for two weeks.

Diffraction data were collected at either the National Synchrotron Light Source II (NSLS-II) beamline 17-ID-2 or the Advanced Photon Source (APS) NE-CAT beamline 24-ID-E. X-ray intensities and data reduction were evaluated and integrated using XDS (*51*) and merged/scaled using Pointless/Aimless in the CCP4 program suite (*52*). Structure determination and refinement starting phases were obtained by molecular replacement using Phaser (*53*) using the designed model for the structures. Following molecular replacement, the models were improved using phenix.autobuild (*54*) with rebuild-in-place set to false, and using simulated annealing. Structures were refined in Phenix (*54*). Model building was performed using COOT (*55*). The final model was evaluated using MolProbity (*56*). Data collection and refinement statistics are recorded in the statistics table below. Atomic coordinates and structure factors reported in this paper have been deposited in the Protein Data Bank (PDB), with accession code 9PVL (for NovoTag_657_-apo) and 11MQ (for NovoTag_657_-holo).

### Eukaryotic cell culture

HeLa Kyoto cells (Cellosaurus, #CVCL_1922) were cultured at 37 °C with 5% CO_2_ in flasks with Dulbecco’s modified Eagle medium (DMEM) GlutaMax (Gibco, #10566016) supplemented with 10% fetal bovine serum (FBS) (Gibco, #A5256701) and 1% penicillin–streptomycin (PenStrep) (Gibco). HeLa cells (ATCC, #CCL-2) were cultured at 37 °C with 5% CO_2_ in flasks with DMEM (Gibco) supplemented with 10% FBS (Hyclone) and 1% PenStrep (Gibco). HEK293T cells (ATCC, #CRL-3216) were cultured at 37 °C with 5% CO_2_ in flasks with DMEM (Gibco) supplemented with 10% FBS (Hyclone) and 1% PenStrep (Gibco).

### Epifluorescence microscopy of individual NovoTags in living cells

For transient transfection, HeLa CCL-2 cells were seeded at 1.2 × 10^4^ cells per well in an 18-well glass-bottom µ-slide (Ibidi, #81817) and allowed to adhere and grow overnight. 24 h after plating, each well was transfected with 100 ng of plasmid using JetPrime (Polyplus, #101000027), according to the manufacturer’s protocol. NovoTags were targeted to the outer mitochondrial membrane using a mitochondrial localization signal “MitoTag” derived from Tom70 with the amino acid sequence KSFITRNKTAILATVAATGTAIGAYYYYNQLQQ. Cells were transfected to express MitoTag-NovoTag_657_-eGFP, MitoTag-NovoTag_596_-eGFP, MitoTag-NovoTag_494_-mScarlet, MitoTag-NovoTag_657_^cv^-eGFP, MitoTag-NovoTag_657_^cv^, or MitoTag-NovoTag_657_^cv^-eGFP. 16 h after transfection, the culture medium was exchanged for phenol-red-free medium supplemented with the indicated concentrations of 5 nM on target JF dye, and cells were incubated at 37 °C for 15 min to permit labeling. Wells incubated with dyes were then washed three times with fresh phenol-red-free medium; the final wash was left in place for imaging. Fluorescence images were acquired immediately thereafter under live-cell conditions with the OMX-SR microscope (Leica Microsystems). Toptica diode lasers with excitations of 488 nm, 568 nm, or 640 nm were used. Emission was collected using the PCO.edge sCMOS camera using an Olympus 60× 1.42 NA PlanApochromat oil immersion objective; 1024 × 1024 images (pixel size, 6.5 μm) were captured with no binning. Acquisition was controlled with AcquireSR Acquisition control software. Images were deconvolved in SoftWoRx 7.0.0 (GE Healthcare) using the enhanced ratio method and 200 nm noise filtering, followed by background correction with the rolling-ball algorithm in ImageJ/Fiji (sigma = 35). Colocalization analysis was performed using the JaCoP plugin in ImageJ/Fiji (*57*, *58*).

### Epifluorescence imaging of individual NovoTags in fixed cells

Samples were prepared as described above (‘Epifluorescence microscopy of individual NovoTags in living cells’), but cells were only transfected with MitoTag-NovoTag_657_-eGFP. At 16 h post-transfection, replicate wells expressing the same construct were divided into two treatment groups: one group was first fixed with 4% paraformaldehyde (PFA) in dPBS for 15 min at room temperature, washed three times with dPBS, and then incubated with 5 nM JF_657_ for 15 min at 37 °C; the other group was first incubated with 5 nM JF_657_ for 15 min at 37 °C and subsequently fixed under the same conditions. All wells were then washed three times with fresh phenol-red-free medium; the final wash was left in place for imaging. Images were acquired as described above (‘Epifluorescence microscopy of individual NovoTags in living cells’).

### Photostability analysis of NovoTag_657_ in cells

For transient transfection, HeLa Kyoto cells were seeded at 2 × 10^4^ cells per well in an 8-well glass-bottom µ-slide (Ibidi, #80827) and allowed to adhere. 4 h after plating, each well was transfected with 250 ng of plasmid. MitoTag-NovoTag_657_ and MitoTag-HaloTag7 were transfected, and after 24 h, cells were fluorescently labeled with either 50 nM JF_657_ or 50 nM JFX_650_ HaloTag Ligand (Promega, #HT1070) for 30 min at 37 °C, followed by three washing steps with complete growth media, a 15 min incubation at 37 °C, and three additional washing steps. Cells were fixed with 4% PFA in PBS for 15 min at room temperature and washed 3 times with PBS. Fluorescence images were acquired on a confocal microscope LSM900 (Leica Microsystems), equipped with a 40×, 1.20 NA water C-apochromat korr objective and a GaAsP-Pmt2 detector. The intensity of the 640 nm laser was measured using a laser power meter StarLite (Ophir Optronics Solutions) at the sample plane with an inserted objective, yielding 0.743 mW at 100% laser power. For each cell, a central 10 × 10 µm square was selected for photobleaching. Photobleaching was performed with 100% 640 nm laser power, a pixel dwell time of 6.98 µs, and a pixel size of 0.13 µm. Photobleaching followed by acquisition of low-laser-power images was performed for 100 rounds. Image drift was corrected using the ‘correct 3D drift’ implementation in ImageJ/FIJI. Regions of interest within the bleaching area were selected, gray values were extracted, and plotted against the number of bleaching iterations. A nonlinear regression (‘one phase decay’: Y=(Y0 - plateau)*exp(-K*X) + plateau) was fitted using GraphPad Prism (GraphPad Software), and the half-life was determined.

### Evaluation of NovoTag binding specificity in living cells using epifluorescence microscopy

Samples were prepared as described above (‘Epifluorescence microscopy of individual NovoTags in living cells’). Cells were transfected with either 2×FYVE-NovoTag_494_, MitoTag-NovoTag_596_, or H2B-NovoTag_657_. 16 h after transfection, the culture medium was exchanged for phenol-red-free medium supplemented with 5 nM of either JF_494_, JF_596_, or JF_657_, and cells were incubated at 37 °C for 30 min to permit labeling. Wells incubated with dyes were then washed three times with fresh phenol-red-free medium; the final wash was left in place for imaging. Images were acquired using the widefield microscope IN Cell Analyzer 2500HS (Molecular Devices) equipped with a 20× 0.75 NA CFI Plan Apo objective (Nikon). Cells were illuminated with a seven-color Solid State Illuminator (SSI) for fluorescence excitation. Fluorescence signals were acquired sequentially using the following filters: Green (excitation 473 nm, emission 511/23 nm), red (excitation 575 nm, emission 623/24 nm), and far-red (excitation 631 nm, emission 684/24 nm). Imaging was controlled using the IN Cell Analyzer 2500 HS software version 7.4, and light was collected on a sCMOS camera without binning. Evaluation of nonspecific background fluorescence intensity was performed by first background correcting images (Fiji rolling ball algorithm, sigma = 50). Because many negative conditions did not contain signals with which to segment cells, we normalized the image intensities by dividing the mean of all pixel intensities by the 50th percentile pixel intensity. The 50th percentile intensity was confirmed to be consistent with the background pixel intensity, given our cell density and imaging/staining conditions.

### Multiplex epifluorescence microscopy of NovoTags in living cells

Samples were prepared as described above (‘Epifluorescence microscopy of individual NovoTags in living cells’), but cells were co-transfected with MitoTag-NovoTag_657_, Emerin-NovoTag_596_, and 2×FYVE-NovoTag_494_. 16 h after transfection, the culture medium was exchanged for phenol-red-free medium supplemented with a cocktail of 5 nM JF_494_, 20 nM JF_596_, and 5 nM JF_494_ dye, and cells were incubated at 37 °C for 15 min to permit labeling. Wells incubated with dyes were then washed three times with fresh phenol-red-free medium; the final wash was left in place for imaging. Images were acquired as described above (‘Epifluorescence microscopy of individual NovoTags in living cells’).

### Multiplex confocal and super-resolution Tau-STED fluorescence microscopy on fixed cells

For transient transfection, HeLa Kyoto cells were seeded at 2 × 10^4^ cells per well in an 8-well glass-bottom µ-slide (Ibidi, #80827) and allowed to adhere. 24 h after plating, cells were co-transfected with 2×FYVE-NovoTag_494_, MitoTag-NovoTag_596_, and H2B-NovoTag_657_. Each well was transfected with 250 ng of plasmid using TransIT-HeLa transfection reagent (Mirus, #MIR2900) according to the manufacturer’s protocol. After 24 h, cells were fluorescently labeled with 50 nM of each dye (JF_494_, JF_596_, and JF_657_) for 30 min at 37 °C, followed by three washes with complete growth media, a 15 min incubation at 37 °C, and three additional washes. Cells were fixed with 4% PFA in PBS for 15 min at room temperature and washed 3 times with PBS. Super-resolution fluorescence images were acquired on a Stellaris 8 STED Falcon (Leica), equipped with an 86×, 1.20 NA water HC PL APO CS2 motCORR STED WHITE objective, a tunable white light laser, and HyD X detectors. The following excitation and emission setups were used for the three channels: NovoTag_494_: 507 nm excitation laser, 589 nm STED depletion laser, and 517-560 nm emission detection window. NovoTag_596_: 593 nm excitation laser, 775 nm STED depletion laser, and 603-630 nm emission detection window. NovoTag_657_: 661 nm excitation laser, 775 nm STED laser, and 671-720 nm emission detection window. STED image acquisition was performed in photon-counting mode with a pixel size of 22 nm, a pixel dwell time of 1.89 µs, and 8-fold line accumulation using LASX (Leica Microsystems). STED data were processed using the Tau-STED Xtend module in LASX. In addition to STED, confocal images were acquired using the same microscope setup, excitation wavelength, and emission detection window but with deactivated STED laser. Line profiles were measured using ImageJ/FIJI.

### Multiplex confocal and super-resolution Tau-STED fluorescence microscopy on live cells

Sample preparation and staining were performed as described above (section ‘Multiplex confocal and super-resolution Tau-STED fluorescence microscopy on fixed cells’). After the last washing step, the cell culture medium was exchanged for live-cell medium (DMEM FluoroBrite (Gibco) supplemented with 10% FBS and 1% P/S). Images were acquired using the same microscope setup described above, but equipped with an environmental chamber set to 37 °C and 5% CO_2_.

### Computational design pipeline for JF_494_ binders with specific fluorescence lifetimes

We reasoned that by modulating the binding site of JF_494_, we can design binders with altered photophysical properties, such as spectra profiles and fluorescence lifetimes. LigandMPNN FastRelax was used to redesign NovoTag_494_ binding pocket. Specifically, residues within 4.0 Å that form critical interactions with JF_494_, particularly aromatic residues, underwent sequence redesign via LigandMPNN and subsequent structural refinement with FastRelax. All remaining residue identities were kept strictly fixed. Similar filtering constraints from the section above (‘Computational design pipeline for dye binder using pseudocycle Cα RFdiffusion’) were applied here. In addition, we filtered to select designs with 4 or more hydrogen bonds and zero π-π stacking towards JF_494_. For further details on the computational design pipeline, see the code repository on GitHub (https://github.com/steffen-klein/novotag-design) or Zenodo (*45*).

### Screening for JF_494_ binders with different lifetimes

Designs were purified following the above described methods in the ‘Small-scale expression and purification of verified binders’ section. The purified proteins were normalized by the molar concentration in a clear flat bottom plate for plate reader (Corning 3881). JF_494_ was added to the plate at a sub-stoichiometric ratio, with a final concentration half that of the protein. JF_494_ concentration, buffer composition, and final assay volume (dictating the optical path length) were kept constant across all wells of the microplate. Based on theoretical derivations and prior protein-dye complexes engineering efforts (*59*, *60*), we reasoned that fluorescence emission intensity provides a practical, albeit semi-quantitative, proxy that positively correlates with fluorescence lifetime. Thus, designed variants exhibiting higher fluorescence emission peaks were expected to yield longer fluorescence lifetimes, and vice versa. To validate the correlation between emission and lifetime dynamics, two designs, each from the highest and lowest emission peak intensities, that still retain high nanomolar binding affinity or better, were selected for *in vitro* photophysical characterization as described above in the ‘Fluorescence spectra scanning’ section. Fluorescence spectra scanning was performed on a Biotek Synergy Neo2 Reader (Agilent). To measure the fluorescence spectra of bound and unbound dyes, we incubated freshly purified JF dye binders with their respective JF dyes at an excess molar ratio of binders (protein) to dyes. After incubation at room temperature for 5 min, followed by an additional 25 min on a vibrating shaker (Heidolph Titramax 1000 Vibrating Shaker) at 1,000 rpm, 25 °C, the mixture was transferred to a Corning 96 well half area flat bottom non-binding surface black plate (Corning, #3686). Subsequently, equal volumes and concentrations of JF dyes (diluted in TBS buffer) were transferred to the plate. For fluorescence emission spectral scanning, samples containing JF_494_ were excited at 505 nm, and emission was scanned from 515 to 620 nm with a 1 nm step size. The binding characterization followed the method described in the ‘Fluorescence polarization (FP) assay for dye binder affinity of binders.’

### Multiplex confocal fluorescence lifetime imaging microscopy (FLIM)

For transient transfection, HeLa Kyoto cells were seeded at 3 × 10^3^ cells per well in an 8-well glass-bottom plate (Ibidi, µ-slide, #80827) and allowed to adhere. 24 h after plating, cells were co-transfected with plasmids encoding for 2×FYVE-NovoTag_494_, MitoTag-NovoTag_494_^S^, and H2B-NovoTag_494_^L^. Each well was transfected with 250 ng of total plasmid using TransIT-HeLa transfection reagent (Mirus, #MIR2900) according to the manufacturer’s protocol. After 24 h, cells were fluorescently labeled with 50 nM JF_494_ for 30 min at 37 °C and chemically fixed with 4% PFA for 15 min at room temperature. Samples were imaged in PBS with 10 nM JF_494_. Fluorescence images were acquired on a confocal microscope Stellaris 8 Falcon FLIM (Leica Microsystems) equipped with a 40× 1.20 NA water HC PL APO CS2 objective, tunable pulsed white light laser (470-670 nm), and a HyD detector. The following excitation and emission setup was used: 507 nm excitation laser and 517-607 nm emission detection window. Images were acquired with a pixel size of 94 nm and a scan speed of 400 Hz using LAS X (Leica Microsystems). Fluorescence lifetime data were analyzed using LAS X FLIM software (Leica Microsystems). Lifetime decays were fitted using a two-component exponential model. Lifetime unmixing was performed with the Phasor module in LAS X FLIM (Leica Microsystems).

### Fluorescence lifetime measurements in cells

To measure the lifetimes of the three different NovoTag_494_ variants (NovoTag_494_, NovoTag_494_^S^, and NovoTag_494_^L^) in cells, samples were prepared as described above (section ‘Multiplex confocal fluorescence lifetime imaging microscopy (FLIM)’), but only one plasmid per sample was transfected. Image acquisition was performed as described above, but with a 200 Hz scanning speed. Fluorescence lifetime data were analyzed using LAS X FLIM software (Leica Microsystems). Lifetime decays were fitted using a two-component exponential model. Regions of interest were selected and lifetime histograms were generated.

### Computational design pipeline for covalent dye binder using an S_N_Ar reaction

The design of the JF_657_ covalent binders was based on 36 experimentally validated non-covalent JF_657_ binders. These binders were first filtered to select structures where at least one residue Cα atom was within 4.0 Å of the para-fluorine from JF_657_. The residues satisfying this geometric criterion were then mutated to cysteine as the designated nucleophile for an attack on the para-fluorine-bearing carbon. We then used Rosetta FastRelax (*13*) to further filter for designs with optimal distance and geometry between the thiol group and para-fluorine. For further details on the computational design pipeline, see the code repository on GitHub (https://github.com/steffen-klein/novotag-design) or Zenodo (*45*).

### S_N_Ar protein identification through fluorescent SDS-PAGE protein gel electrophoresis

To screen for NovoTag_657_^cv^, we first expressed designed proteins as described in ‘Small-scale expression and purification of verified binders’. These proteins were then purified on HPLC as described above (‘Small-scale expression and purification of verified binders’) . Proteins at the expected peaks were incubated with 10 µM JF_657_ at 37 °C in 20 mM Tris, 100 mM NaCl, pH 8 (TBS), for 2 and 12 h. The sample was incubated in a 1.5 ml microcentrifuge tube (Eppendorf™ #05-402-27) at 37 °C with shaking at 1,000 rpm on a ThermoMixer (Eppendorf, #05-412-503). The samples were prepared for SDS-PAGE by adding LICOR 4X Protein Sample Loading Buffer (LICOR 92840004) to the incubated protein-JF_657_ mixture, and the mixture was heated at 95 °C for 5 min, then loaded onto Mini-PROTEAN TGX Stain-Free Precast AnykD Gels (Bio-Rad, #456-8126). Protein ladder (Precision Plus Protein Dual Color Standards #1610374 or Chameleon Duo Pre-stained Protein Ladder #928-60000) was added. Gel electrophoresis was performed at 200 V for 30 min. The gel was then imaged using the LI-COR Odyssey M imager (excitation 685 nm, emission 721-740 nm) to identify the far-red fluorescent signal. Afterward, the gel was stained with Coomassie blue. LI-COR Odyssey M imager (excitation 631-659 nm, emission clear window) was used to identify the protein bands.

### S_N_Ar covalent labeling analysis in *E. coli* through fluorescent SDS-PAGE protein gel electrophoresis

To validate NovoTag_657_^cv^ covalent labeling in living cells, *E. coli* BL21 cells expressing NovoTag_657_^cv^ were cultured in Terrific Broth TB auto-induction medium supplemented with 1 µM of JF_657_, at 37 °C. After 12 or 18 hours, the cultures were harvested. The resulting cell pellets were washed three times with TBS to remove residual JF_657_. Finally, cells were lysed, and the whole-cell lysate was analyzed by SDS-PAGE as described above (‘S_N_Ar protein identification through fluorescent SDS-PAGE protein gel electrophoresis’).

### S_N_Ar covalent labeling analysis in HeLa cells through fluorescent SDS-PAGE protein gel electrophoresis

HeLa CCL-2 cells were seeded at 1.2 × 10^4^ cells per well in an 18-well glass-bottom µ-slide (Ibidi, #81817) and allowed to adhere and grow overnight. 24 h after plating, each well was transfected with 100 ng of the plasmid encoding MitoTag-NovoTag_657_^cv^ using JetPrime (Polyplus, #101000027), according to the manufacturer’s protocol. Additionally, 1 µM JF_657_ was supplemented in the DMEM medium. After 18 h, the cells were washed three times with cold dPBS to remove residual JF_657_. Cells were subsequently lysed for 15 min in RIPA buffer (Thermo Fisher, #89900) supplemented with Halt Protease and Phosphatase Inhibitor Cocktail (Thermo Fisher, #87785). The resulting whole-cell lysates were then analyzed by SDS-PAGE.

### Liquid chromatography-mass spectrometry (LC-MS) for S_N_Ar labeling kinetics evaluation

Protein samples after size-exclusion chromatography purification were normalized to 10 μM (about 0.18 mg/ml) and subsequently incubated with 50 μM JF_657_. Both proteins and JF_657_ were prepared in TBS (20 mM Tris, 100 mM NaCl, pH 8). The sample was incubated in a 1.5 ml microcentrifuge tube (Eppendorf, #05-402-27) at 37 °C with shaking at 1,000 rpm on a ThermoMixer (Eppendorf, #05-412-503). After a specific incubation period (0.5 h, 1 h, 2 h, 3 h, and 12 h), the incubated samples were buffer-exchanged into TBS using Zeba Spin Desalting Columns (Thermo Fisher Scientific) to remove excess JF_657_. LC-MS was obtained under positive mode on an Agilent G6230B Time-of-Flight (ToF) LC-MS equipped with an AdvanceBio RP-Desalting column, and subsequently deconvoluted by using a total entropy algorithm from Agilent MassHunter BioConfirm software. The integrated peak area (mass area) of both labeled and non-labeled species was extracted to calculate labeling efficiency.

### Computational design pipeline for a chemically induced dimerization system

The design of a chemically induced dimerization system was based on the top 11 experimentally validated non-covalent JF_657_ binder designs. For each design, all helix-loop-helix positions were identified. Next, all possible split variants based on those positions were generated. To that aim, neighboring helices were grouped into two groups, and the design was split at one or two helix-loop-helix regions, and C-and N-termini were fused with a flexible linker if necessary. The minimum group size was set to 2 helices. In total, 132 split variants were generated. These designs were filtered for high confident protein-small molecule complexes formation, as reflected by RoseTTAFold-All-Atom (*61*) protein-ligand-PAE score (<= 13), AF2 multimer (*62*) pLDDT score (>= 92), and US-align (*63*) TMscore between design and AF2 prediction (>= 0.92). For further details on the computational design pipeline, see the code repository on GitHub (https://github.com/steffen-klein/novotag-design) or Zenodo (*45*).

### Mammalian two-hybrid screen for a chemically induced dimerization system

To analyze the JF_657_-induced dimerization of the split designs, an adapted mammalian two-hybrid screen (*64*, *65*) was used, which is based on the CheckMate system (Promega): The two components of the split system were cloned into two plasmids that contain either a ZF6 or p65 cassette. In addition, a reporter plasmid was used that contains an 8×ZF6 binding site, which controls expression of mNeonGreen. Thus, mNeonGreen expression is a readout of dimerization of the split design. For the mammalian two-hybrid screen, 1 × 10^4^ HEK293T cells per well were seeded in a 24-well plate (Nunc, Thermo Fisher Scientific). After 24 h, cells were transfected with the two plasmids for each split monomer and the reporter plasmid using PEI. 24 h post-transfection, cells were treated with 500 nM JF_657_. The control group was not treated with JF_657_. After 24 h, cells were trypsinized and prepared for FACS analysis using Attune CytPix Flow Cytometer. mNeonGreen fluorescence signal was gated using the laser with excitation at 488 nm and emission at 585 nm. Samples without a fluorescence signal in the non-induced state, but with a fluorescence signal in the induced state, were selected for microscopy-based colocalization validation.

### Epifluorescence microscopy colocalization screen for a chemically induced dimerization system

For each split design, a plasmid was synthesized, encoding a MitoTag, followed by JF_657_ binding protein (split A), mScarlet, then ribosomal skipping P2A, followed by JF_657_ binding protein (split B), fused to mNeonGreen. Thus, JF_657_ binding protein (split A) is localized at the outer mitochondrial membrane, whereas JF_657_ binding protein (split B) is freely expressed in the cytosol. Induced dimerization of the split system should lead to colocalization of mScarlet and mNeonGreen. For transient transfection, HeLa Kyoto cells were seeded at 8 × 10^3^ cells per well in an 8-well glass-bottom µ-slide (Ibidi, #80827) and allowed to adhere. After 2 h, cells were transfected with 150 ng plasmid per well using Lipofectamine LTX (Thermo Fisher Scientific) according to the manufacturer’s protocol. After 24 h, the experimental group was treated with 200 nM JF_657_ for 1 h at 37 °C. Cells were fixed with 4% PFA in PBS for 15 min at room temperature and washed 3 times with PBS. Images were acquired on a widefield fluorescence microscope Ti-Eclipse (Nikon), equipped with a 40×, 0.95 NA air CFI P-Apo objective, a SpectraX illumination (Lumencor), and an Orca Flash 4 V2 camera (Hamamatsu). The following excitation and emission setups were used for the three channels: mNeonGreen: excitation filter: 485/20 nm, emission filter: 447/60 nm, mScarlet: excitation filter: 560/25 nm, emission filter: 593/40 nm, and for JF_657_: excitation filter: 650/13 nm, emission filter: 692/40 nm. Z-stacks were acquired using a Z-step size of 1 µm and a pixel size of 162.5 nm. Data was deconvolved using Huygens Professional (Scientific Volume Imaging). Colocalization analysis between mScarlet and mNeonGreen was performed using the JaCoP plugin in ImageJ/Fiji (*57*, *58*).

### Structure-guided site-directed mutagenesis of NovoSplit_657_ to minimize non-induced self-dimerization

The top design of the colocalization screen described above was selected, and the structure was predicted using AF2 multimer. Potential salt bridges between the split monomers were identified using PyMOL. For each identified salt bridge, one side chain was replaced with alanine to prevent its potential formation. Colocalization analysis was used to evaluate different amino acid substitutions: For transient transfection, HeLa Kyoto cells were seeded at 2 × 10^4^ cells per well in an 8-well glass-bottom plate (Ibidi, µ-slide, #80827) and allowed to adhere. 4 h after plating, cells were co-transfected with two plasmids encoding for each monomer: MitoTag-mStayGold-NovoSplit_657_^A^ and MitoTag-mCherry-NovoSplit_657_^B^ (with different amino acid substitutions). Each well was transfected with 250 ng of total plasmid using TransIT-HeLa transfection reagent (Mirus, #MIR2900) according to the manufacturer’s protocol. After 24 h, cells were fluorescently labeled with 200 nM JF_657_ for 60 min at 37 °C. Cells were fixed with 4% PFA in PBS for 15 min at room temperature and washed 3 times with PBS. Fluorescence images were acquired on a confocal microscope LSM 980 AIRY (Zeiss) equipped with a 40×, 1.20 NA water C-Apochromat autocorr FCS M27 objective, and a spectral detector. The following excitation and emission setups were used for the three channels: mStayGold: 488 nm excitation laser and 495-550 nm emission filter. mCherry: 561 nm excitation laser and 573-627 nm emission filter. JF_657_: 639 nm excitation laser and 300-735 nm emission filter. Images were acquired in frame-sequential mode with a pixel size of 99 nm and a pixel dwell time of 1.67 µs using ZEN Blue (Zeiss). AiryScan images were processed with identical settings across all datasets using Zen Blue (Zeiss). Regions of interest for individual cells were manually selected, and colocalization analysis between mCherry, mStayGold, and JF_657_ was performed using the coloc2 plugin in ImageJ/Fiji (*58*). For each dataset, between 28 and 99 individual cells were analyzed.

### Confocal fluorescence live cell microscopy for NovoSplit_657_

For transient transfection, HeLa Kyoto cells were seeded at 5 × 10^3^ cells per well in an 8-well glass-bottom plate (Ibidi, µ-slide, #80827) and allowed to adhere. 4 h after plating, cells were co-transfected with two plasmids encoding for each monomer: MitoTag-mStayGold-NovoSplit_657_^A^ and MitoTag-mCherry-NovoSplit_657_^B^. Each well was transfected with 250 ng of total plasmid using TransIT-HeLa transfection reagent (Mirus, #MIR2900) according to the manufacturer’s protocol. After 24 h, culture medium was exchanged with live-cell medium (DMEM FluoroBrite (Gibco) supplemented with 10% FBS and 1% P/S). Fluorescence images were acquired on a confocal microscope LSM 980 (Zeiss) equipped with a 40× 1.20 NA water C-Apochromat autocorr FCS M27 objective, and a GaAsP-PMT detector. The following excitation and emission setups were used for the three channels: mStayGold: 488 nm excitation laser, and 499-552 nm emission detection window. mCherry: 561 nm excitation laser and 570-641 nm emission detection window. JF_657_: 639 nm excitation laser and 650-694 nm emission detection window. Images were acquired in frame-sequential mode with a pixel size of 99 nm and a pixel dwell time of 0.736 µs using ZEN Blue (Zeiss). During imaging, an environmental chamber was set to 37 °C and 5% CO_2,_ and a Define Focus System (Zeiss) was utilized to keep the same focal plane in focus during imaging. Every 3 min a frame was acquired. After 12 min of image acquisition, live cell culture medium was exchanged with fresh live cell culture medium containing 1 µM JF_657_. Live cell imaging was performed for a total of 60 min. Regions of interest for individual cells were selected for each time point, and colocalization analysis between mCherry and mStayGold was performed using the coloc2 plugin in ImageJ/Fiji (*57*, *58*). Pearson’s colocalization coefficients were plotted against time, and a nonlinear regression (plateau followed by one phase association) (Y= IF( X<X0, Y0,Y0 + (Plateau-Y0)*(1 - exp(-K*(X-X0))))) was fitted and the half times (t_1/2_=(1/k)*ln((Y0+YM)/Y0)) were determined with GraphPad Prism.

### Epifluorescence microscopy analysis of NovoSplit_657_ as a proximity biosensor

Cell culture and transfection were performed as described in the section above (‘Epifluorescence microscopy of individual NovoTags in living cells’), using two groups of co-transfection: experimental group was co-transfected plasmids encoding MitoTag-LHD(A)-NovoSplit_657_^A^-mScarlet and LHD(B)-NovoSplit_657_^B^-mNeonGreen, and the control group was co-transfected with MitoTag-LHD(A)-NovoSplit_657_^A^-mScarlet and NovoSplit_657_^B^-mNeonGreen. At 16 h post transfection, cells were fixed with 4% PFA in dPBS for 15 min at room temperature and washed 3 times with dPBS. After fixation, cells were treated with 5 nM of JF_657_, diluted in phenol-red-free medium (DMEM FluoroBrite, Gibco), incubated at 37 °C for 30 min to permit labeling. Cells were then washed three times with dPBS, the final wash was replaced with fresh phenol-red-free medium prior to imaging. Images were acquired as described above (‘Epifluorescence microscopy of individual NovoTags in living cells’).

## Supplementary figures

**Fig. S1.**
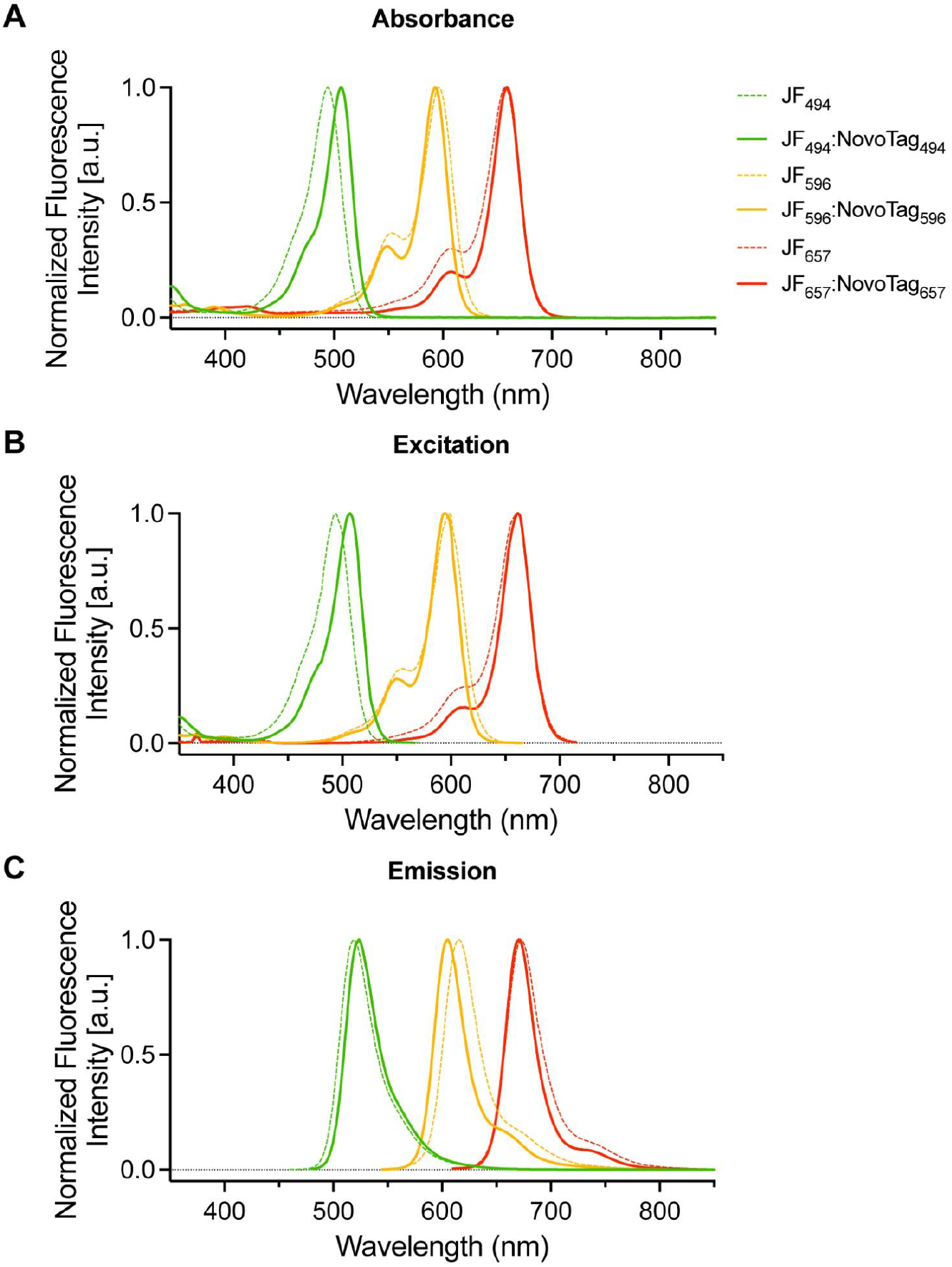
Absorbance, excitation, and emission spectra of JF dyes free in solution or bound to NovoTags. **(A – C)** For each JF dye, JF_494_ (green), JF_596_ (orange), and JF_657_ (red), the absorbance (A), excitation (B), and emission (C) spectra are shown for the free dye (dotted lines) or bound to their respective NovoTags (solid lines).

**Fig. S2.**
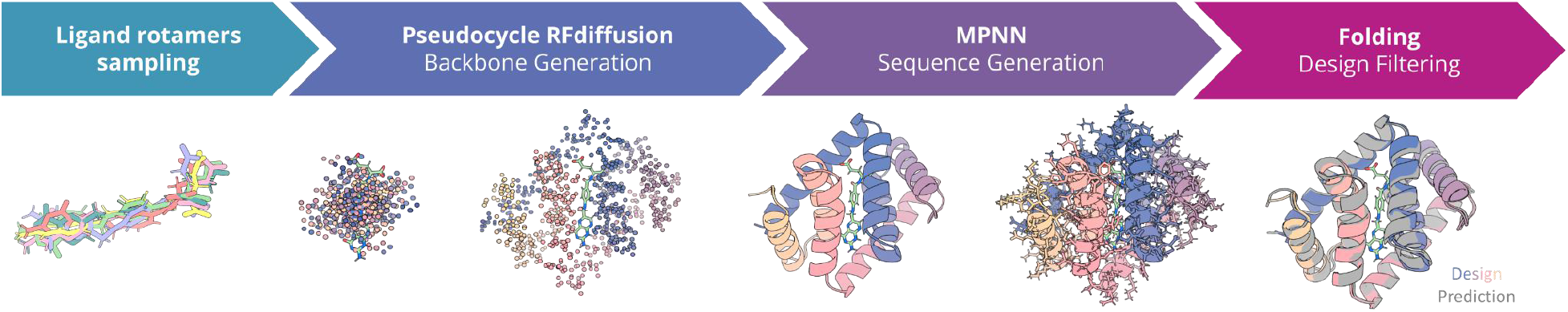
Computational design pipeline. Ligand rotamers were generated using CREST (*27*). Protein backbones were generated using pseudocycle RFdiffusion (*9, 10*). Sequences were generated using LigandMPNN (*12*). Protein folding and ligand binding were assessed using Rosetta (*14*) and AF2 (*15*).

**Fig. S3.**
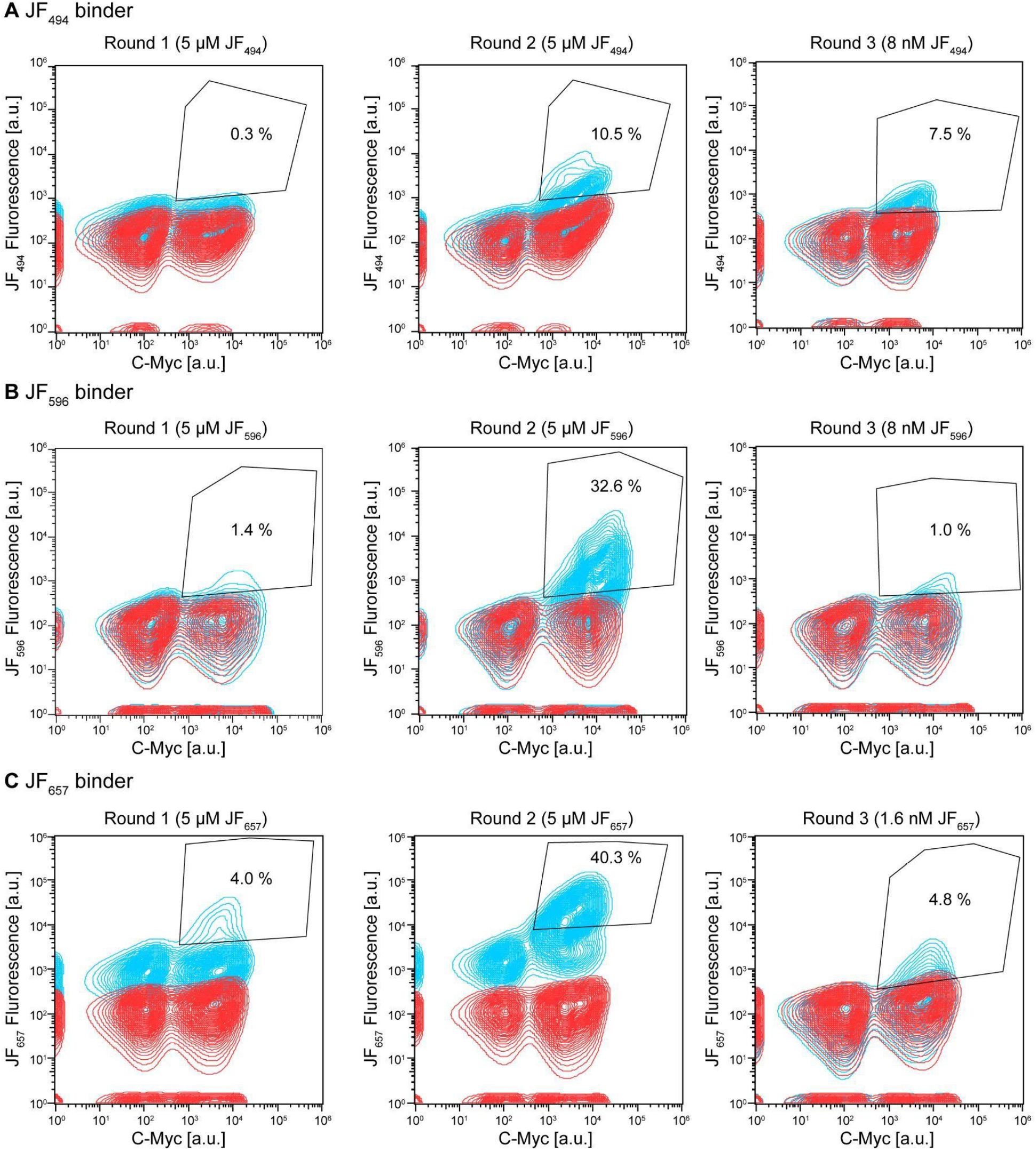
FACS of JF dye binder libraries on yeast surface display. **(A – C)** Each JF dye binder library for JF_494_ (A), JF_596_ (B), and JF_657_ (C), underwent three rounds of binding enrichment via FACS. Enrichment rounds 1 and 2 were performed using 5 μM of on-target JF dye. Round 3 was the titration sorting, performed using 8 nM JF_494_ (A), 8 nM JF_596_ (B), and 1.6 nM JF_657_ (C). The x-axis indicates fluorescence intensity of C-Myc antibody labeling (protein display), while the y-axis indicates fluorescence intensity of the on-target JF dye. Contour plots represent the cells with on-target JF dye incubation (blue) and cells without dye incubation (red). The percentage of cells identified as binding (sorted) over total cell reads is represented within the binding polygon gates.

**Fig. S4:**
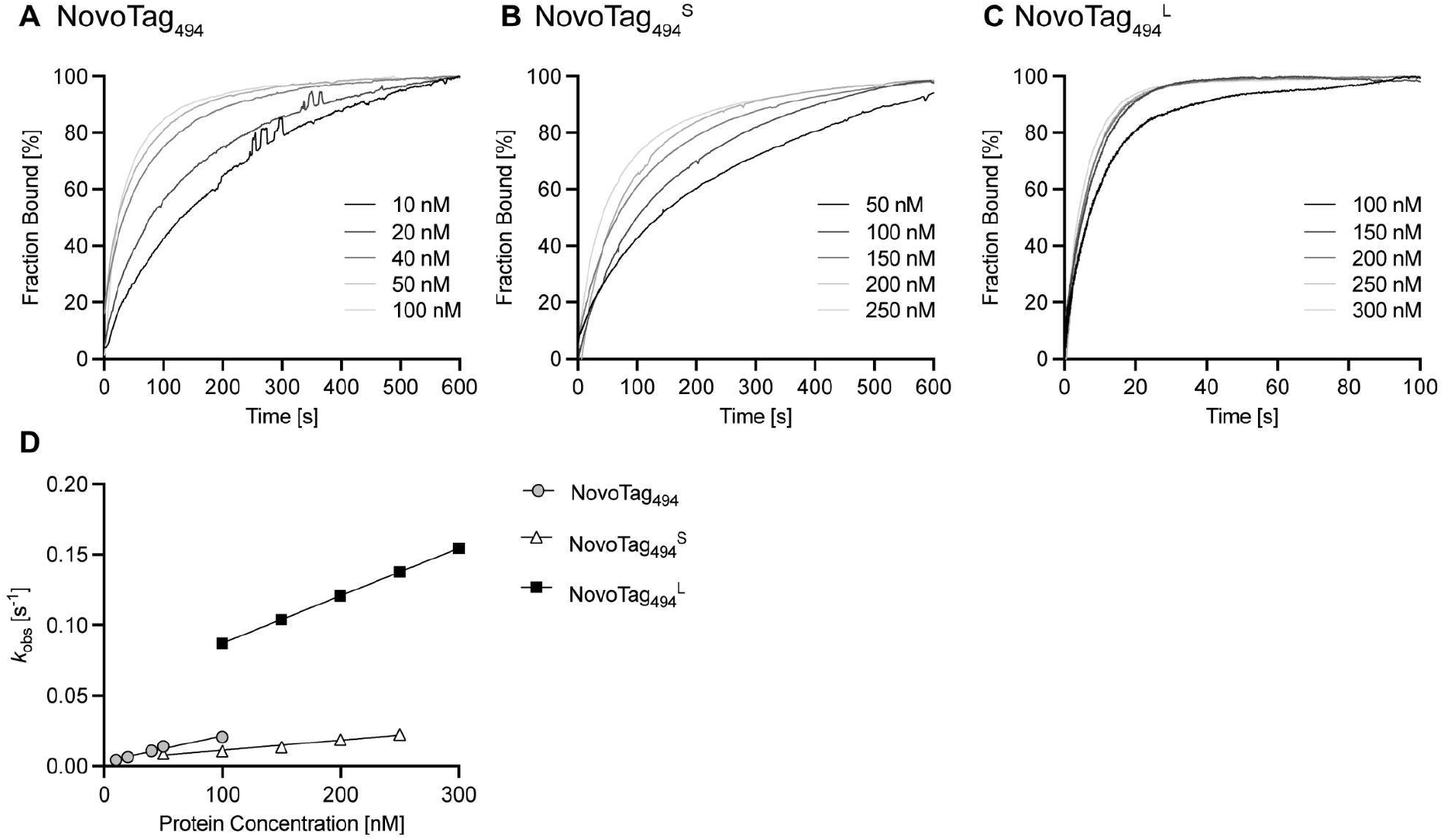
Binding kinetics of JF_494_ to NovoTag_494_ and NovoTag_494_ lifetime variants. Dye binding kinetics were determined using a stopped-flow instrument with fluorescence polarization detection. The percent dye binding is plotted based on the maximum anisotropy for each trace using the average of triplicate measurements. (**A**) Binding curves for NovoTag_494_ with different protein concentrations yielding a k_on_ of 1.8 × 10^5^ M^−1^s^−1^ and k_off_ of 0.00336 s^−1^. (**B**) Binding curves of NovoTag_494_^S^ variant, with different protein concentrations yielding a k_on_ of 7 × 10^4^ M^−1^s^−1^ and k_off_ of 0.0044 s^−1^. (**C**) Binding curves of NovoTag_494_^L^ variant, with different protein concentrations yielding a k_on_ of 3.37 × 10^5^ M^−1^s^−1^ and k_off_ of 0.0044 s^−1^. The data in (A) and (B) display single-phase binding kinetics; the data in (C) display two-phase binding kinetics. The initial fast phase rate is unresolved. Therefore, rates were calculated from the second, slower binding phase. (**D**) Global plots of k_obs_ vs. protein concentrations for the three NovoTag_494_ variants showing the expected linear correlations.

**Fig. S5.**
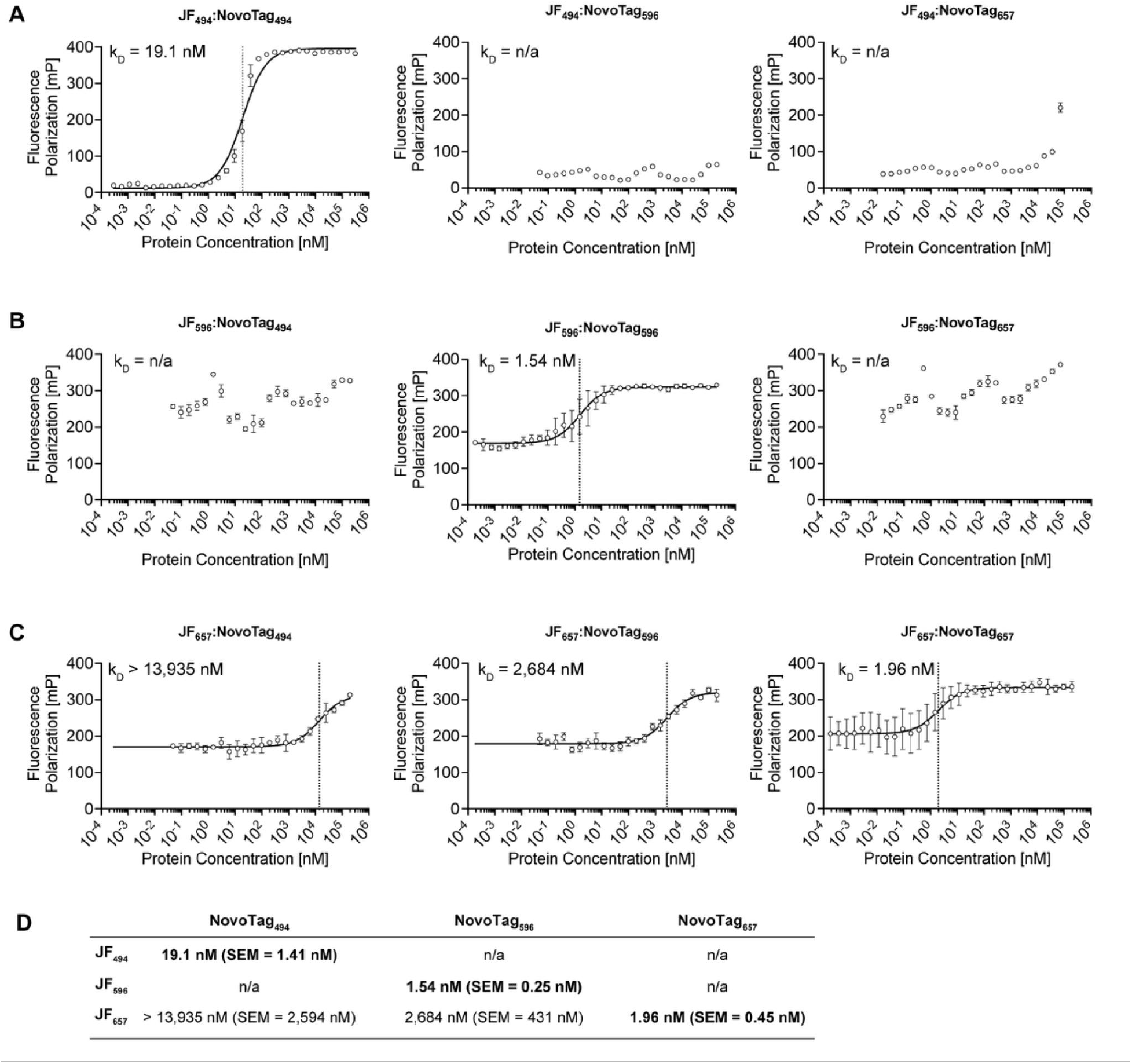
Fluorescence polarization titrations for all combinations of NovoTags and JF dyes. **(A – C)** Fluorescence polarization (FP) titration was acquired for each combination of NovoTags (NovoTag_494_, NovoTag_596_, NovoTag_657_) and JF dyes (JF_494_, JF_596_, JF_657_) as indicated. For each FP titration, 3-8 independent measurements were acquired. For each concentration, the FP (mean ± SD) is shown. A non-linear regression (standard binding isotherm model) was fitted, and K_d_ values were determined. **(D)** Overview of average K_d_ values with indicated SEM.

**Fig. S6.**
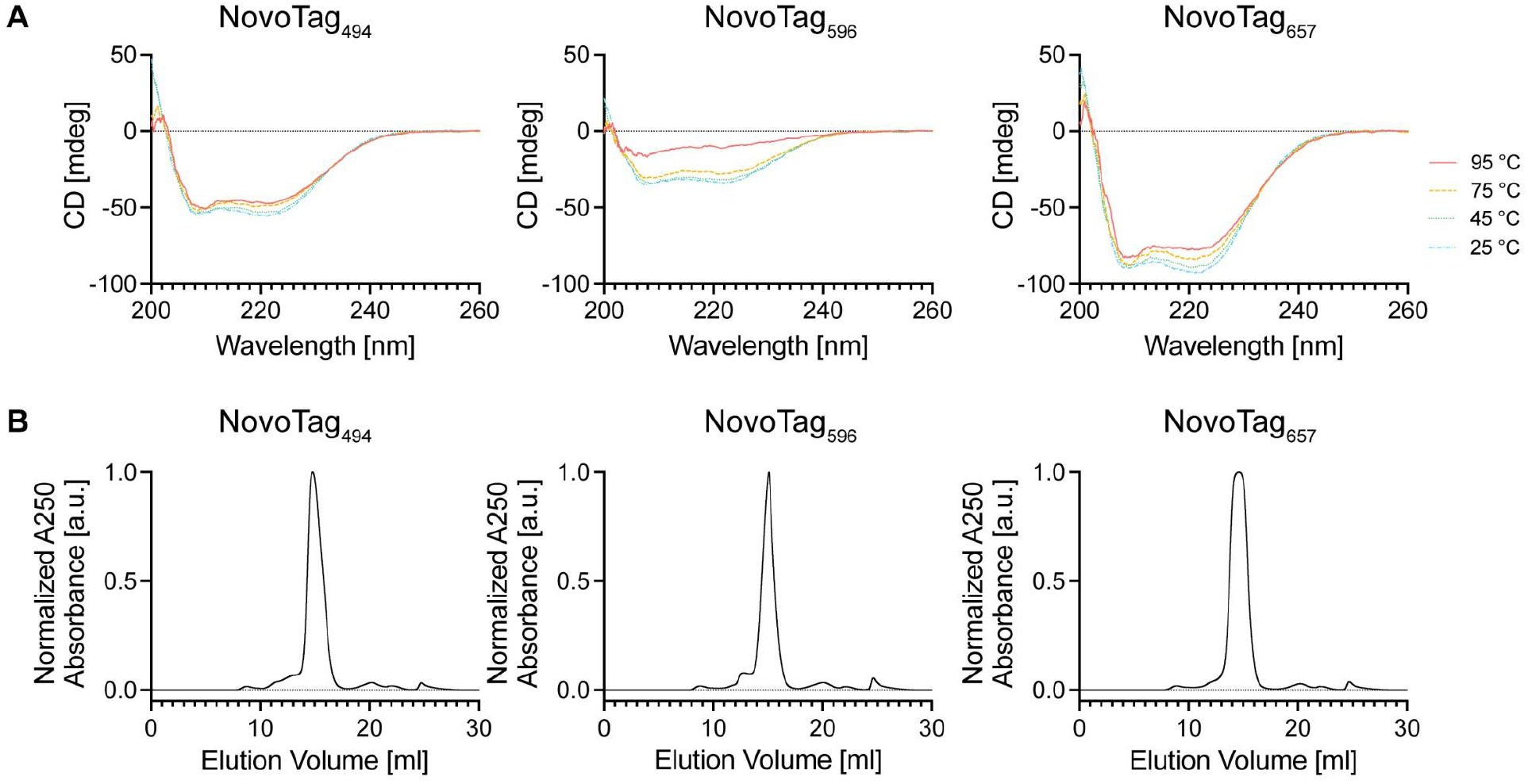
Biochemical characterization of NovoTags. **(A & B)** Circular dichroism analysis of NovoTags at increasing temperatures (A), and size exclusion chromatography analysis of NovoTags, yielding monomeric protein profiles (B).

**Fig. S7.**
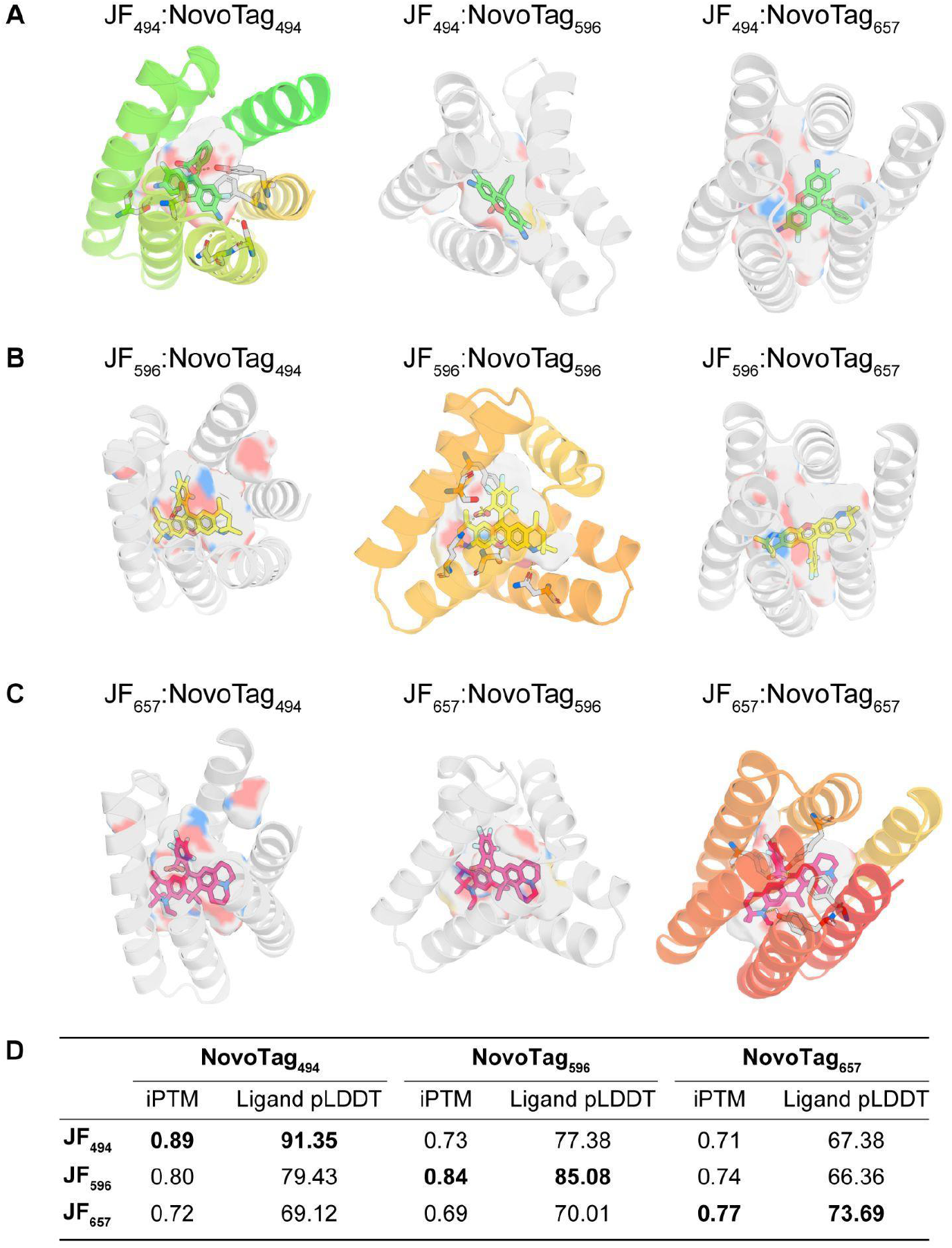
*In silico* structural prediction and dye specificity analysis of NovoTags. **(A – C)** AF3 prediction of the three NovoTags (NovoTag_494_, NovoTag_596_, and NovoTag_657_) co-folded with JF_494_ (A), JF_596_ (B), and JF_657_ (C). **(D)** AF3 metrics (iPTM and ligand pLDDT) for all combinations of NovoTags and dyes. On-target pairs yield higher ipTM and ligand pLDDT (shown in color in A-C) over off-target pairs (shown in grey).

**Fig. S8.**
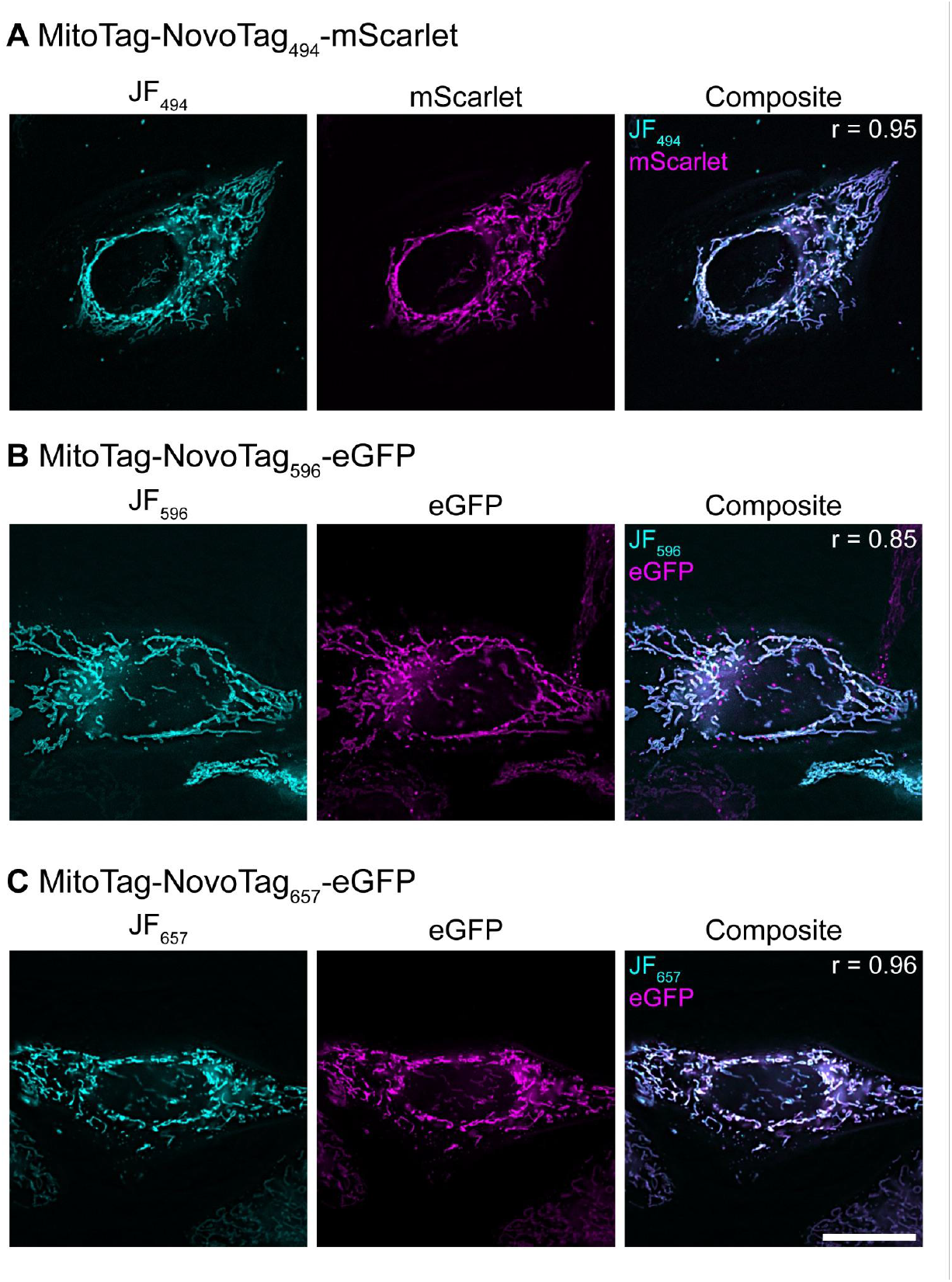
Fluorescence microscopy in live cells using NovoTags. **(A – C)** HeLa cells were transfected with one of the following constructs: MitoTag-NovoTag_494_-mScarlet (A), MitoTag-NovoTag_596_-eGFP (B), or MitoTag-NovoTag_647_-eGFP (C). Cells were incubated with 5 nM of the corresponding JF dye (JF_494_, JF_596_, JF_657_) for 15 min. After three washing steps, live cells were imaged on an OMX SR microscope. For each construct, both fluorescent channels (of the fluorescent protein fusion and the JF dye) and a composite image are shown. Pearson’s correlation coefficient (r) is shown for each image. Scale bar: 20 µm.

**Fig. S9.**
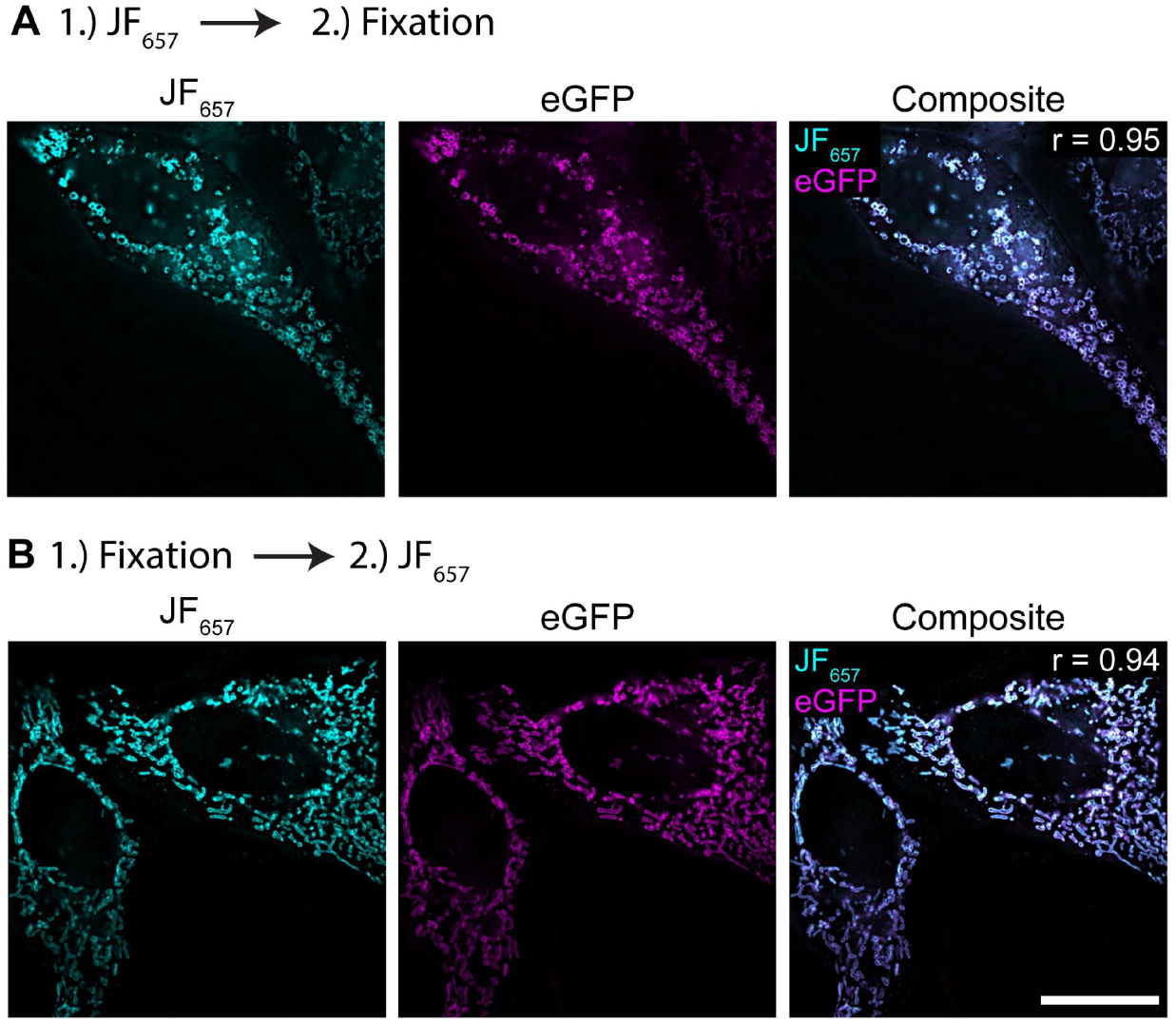
Impact of chemical fixation on NovoTag_657_ for fluorescence microscopy in cells. **(A – B)** HeLa cells were transfected with MitoTag-NovoTag_657_-eGFP. Cells were either first incubated with 5 nM JF_657_ for 15 min at 37 °C, followed by chemical fixation (A), or first chemically fixed, and then incubated with 5 nM JF_657_ for 15 min (B). After three washing steps, cells were imaged on an OMX SR microscope. For each sample, both fluorescent channels (of the fluorescent protein fusion and the JF dye) and a composite image are shown. Pearson’s correlation coefficient (r) is shown for each image. Scale bar: 20 µm.

**Fig. S10.**
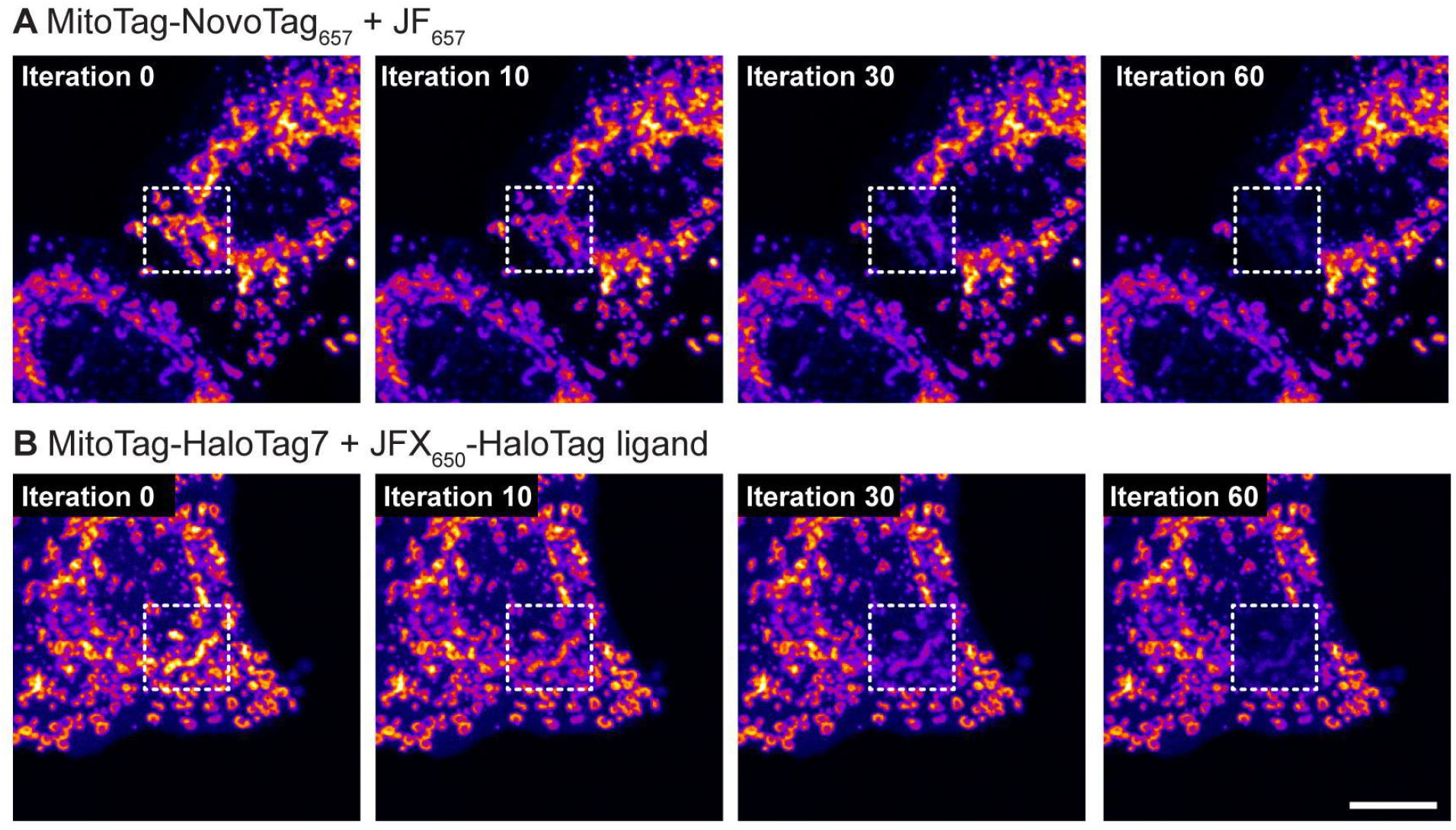
Photostability comparison of the designed NovoTag_657_ to HaloTag7. **(A & B)** HeLa cells were transfected with MitoTag-NovoTag_657_ and fluorescently labeled with JF_657_ (50 nM for 30 min) (A), or with MitoTag-HaloTag7 and fluorescently labeled with Halo-JFX_650_ (50 nM for 30 min) (B). After chemical fixation and three washing steps, cells were photobleached (0.74 mW 640 nm laser, 6.89 µs pixel dwell time, 0.13 µm pixel size) for 100 iterations. Four time points (0, 10, 30, and 60 bleaching iterations) are shown. The white squares (dotted lines) indicate the bleaching area. A lookup table was used to represent fluorescent intensities (“Fire”-LUT in ImageJ). See quantification of intensities over time in Fig. 2C. See also **Movie S1**. Scale bar: 10 µm.

**Fig. S11.**
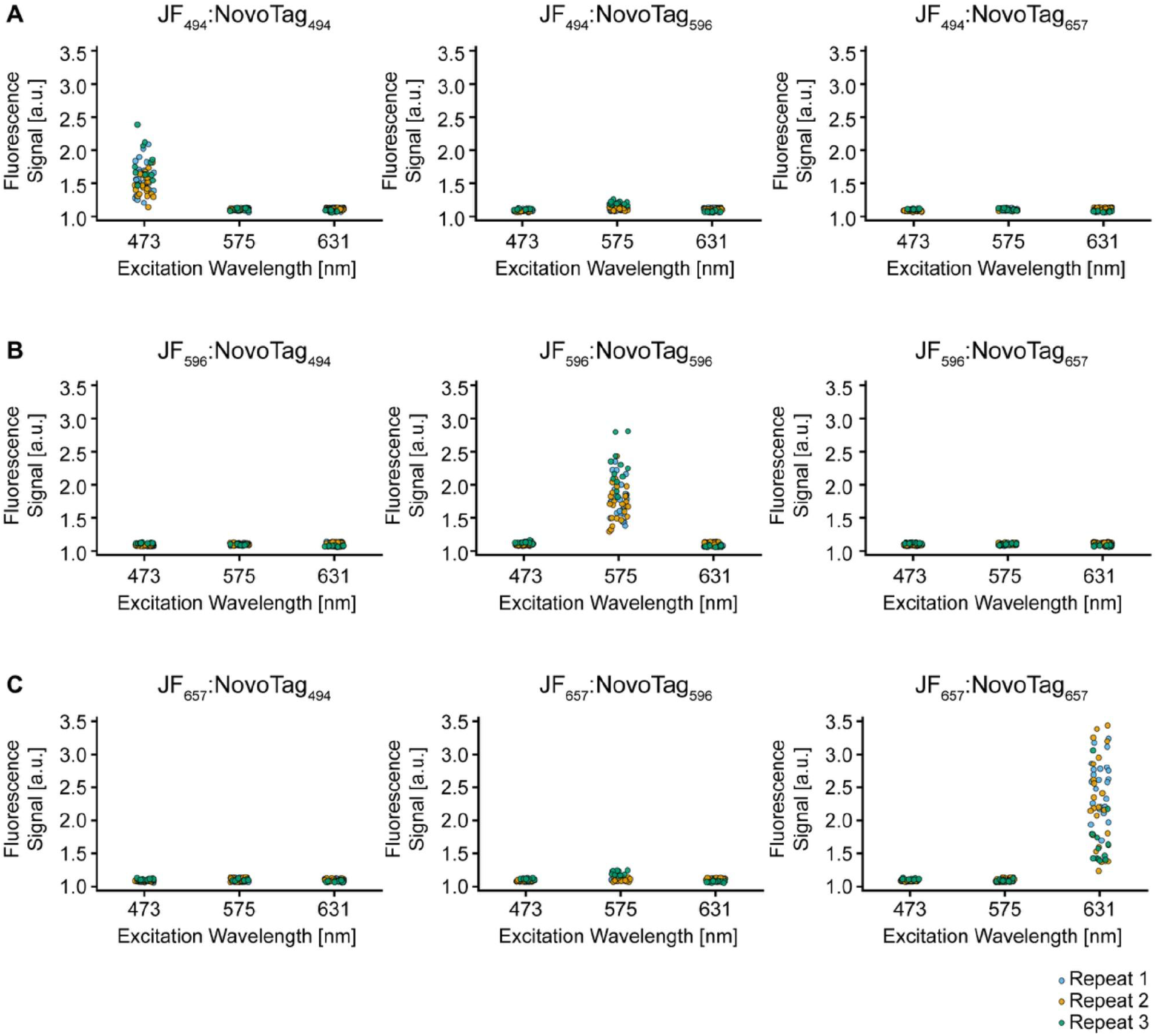
NovoTag dye-binding specificity. **(A – C)** HeLa cells were transfected with 2×FYVE-NovoTag_494,_ H2B-NovoTag_657_, or MitoTag-NovoTag_596_ and fluorescently labeled with the indicated JF dye (JF_494_, JF_596_, or JF_657_) (5 nM for 30 min). Fluorescence images were acquired on the IN Cell Analyzer using the indicated excitation wavelengths (473 nm, 575 nm, 631 nm). The normalized fluorescence signal of each field of view is plotted. For each condition, 12 fields of view were analyzed from three independent experiments (Repeat 1-3).

**Fig. S12.**
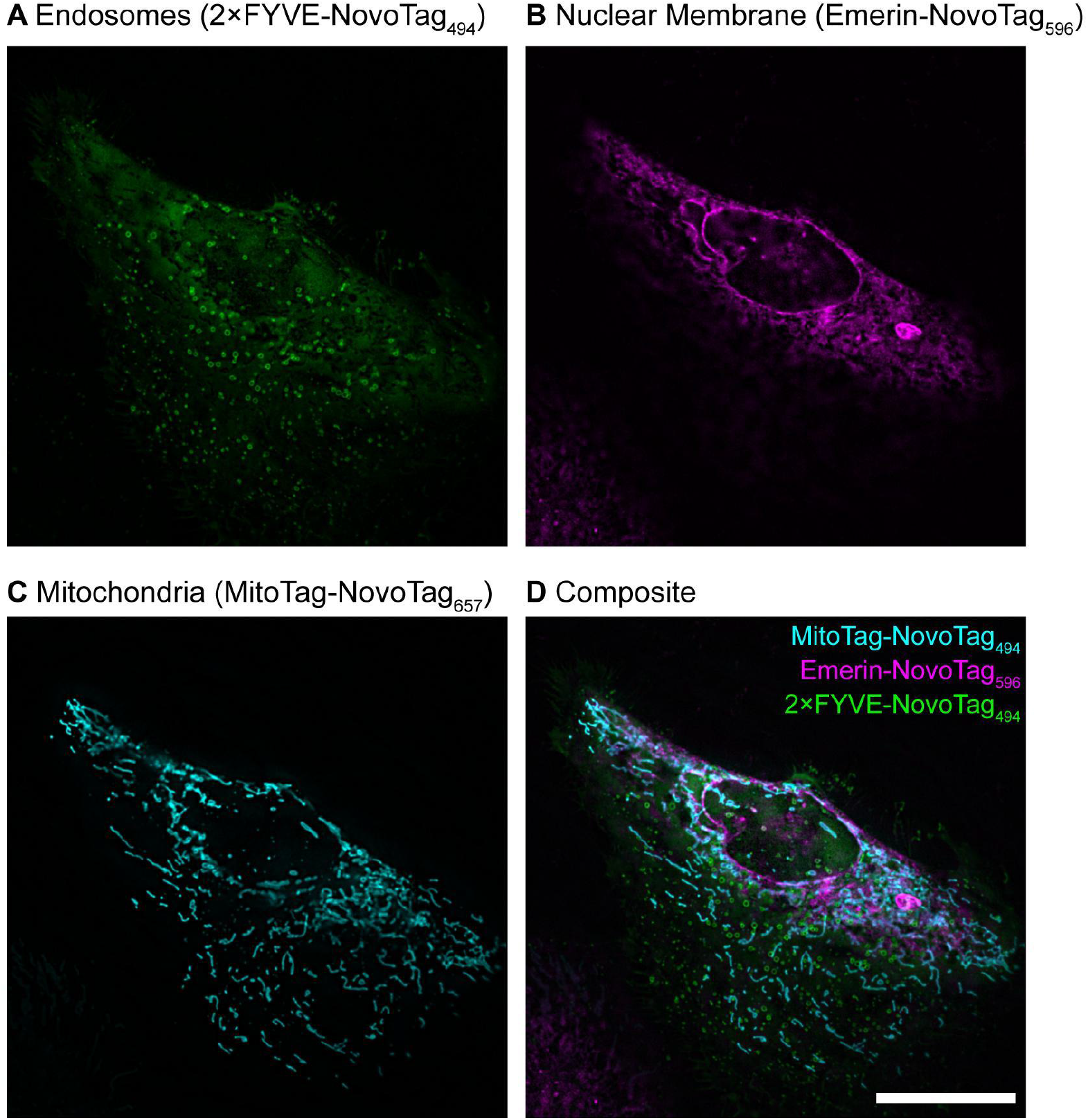
Multiplexed fluorescence microscopy in live cells enabled by orthogonal NovoTags. **(A – D)** HeLa cells were co-transfected with three constructs, targeting different cellular organelles: 2×FYVE-NovoTag_494_ targeting endosomes (A, green), Emerin-NovoTag_596_ targeting the inner nuclear membrane (B, magenta), and MitoTag-NovoTag_657_ targeting the outer mitochondrial membrane (C, cyan). After co-transfection, cells were stained with 5 nM JF_494_, 20 nM JF_596_, and 5 nM JF_657_ for 15 min. After three washing steps, fluorescence microscopy images of live cells were acquired on an OMX SR microscope. Each channel is shown individually (A – C) and as a composite image (D). Scale bar: 20 µm.

**Fig. S13.**
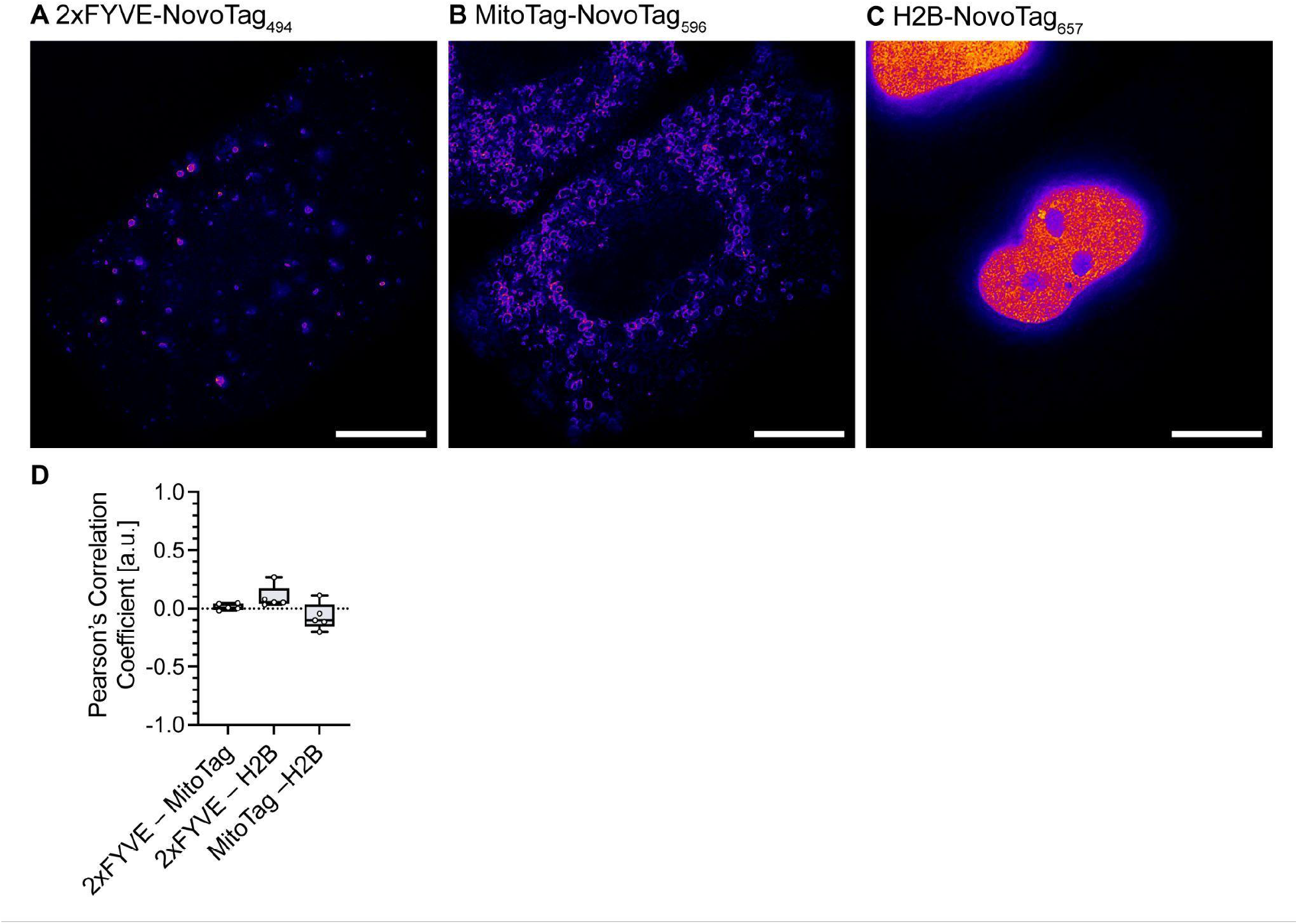
Signal separation of multiplexed STED fluorescence microscopy. **(A – C)** Individual channels of the composite STED image shown in **Fig 2A**. NovoTags were targeted to distinct cellular compartments in HeLa cells: NovoTag_494_ to endosomes (2×FYVE) (A), NovoTag_596_ to mitochondria (MitoTag) (B), and NovoTag_657_ to chromatin (H2B) (C). Cells were stained with 50 nM of each dye (JF_494_, JF_596_, JF_657_) for 30 min at 37 °C, followed by three washing steps. After chemical fixation, STED images were acquired. **(D)** Colocalization between the three labeled components was analyzed. In total, five individual cells were analyzed (n = 5). Scale bar: 10 µm.

**Fig. S14:**
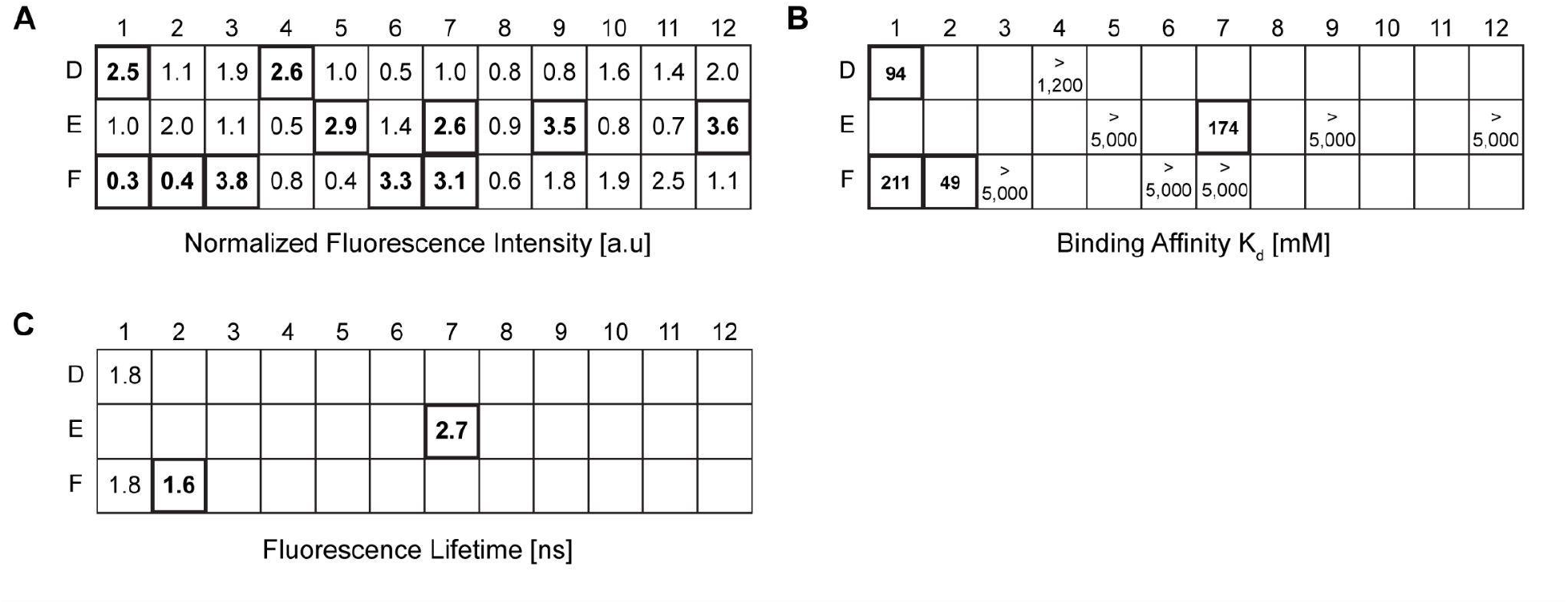
Screening of JF_494_ binder redesigns for fluorescence lifetime variants. **(A)** Normalized peak fluorescence emission intensities of 36 redesigned JF_494_ binders, relative to NovoTag_494_, measured using a BioTek Synergy Neo2 plate reader. The designs with the highest (D1, D4, E5, E7, E9, E12, F3, F6, F7) and lowest (F1, F2) fluorescence intensities were selected for further characterization. **(B)** For selected designs in (A), the binding affinities K_d_ were determined using fluorescence polarization. Designs with the highest binding affinities (D1, E7, F1, F2) were selected for further characterization. **(C)** For selected designs in (B), the *in vitro* fluorescence lifetimes were determined. The design with the shortest (F2, NovoTag_494_^S^) and longest (E7, NovoTag_494_^L^) fluorescence lifetime was selected.

**Fig. S15.**
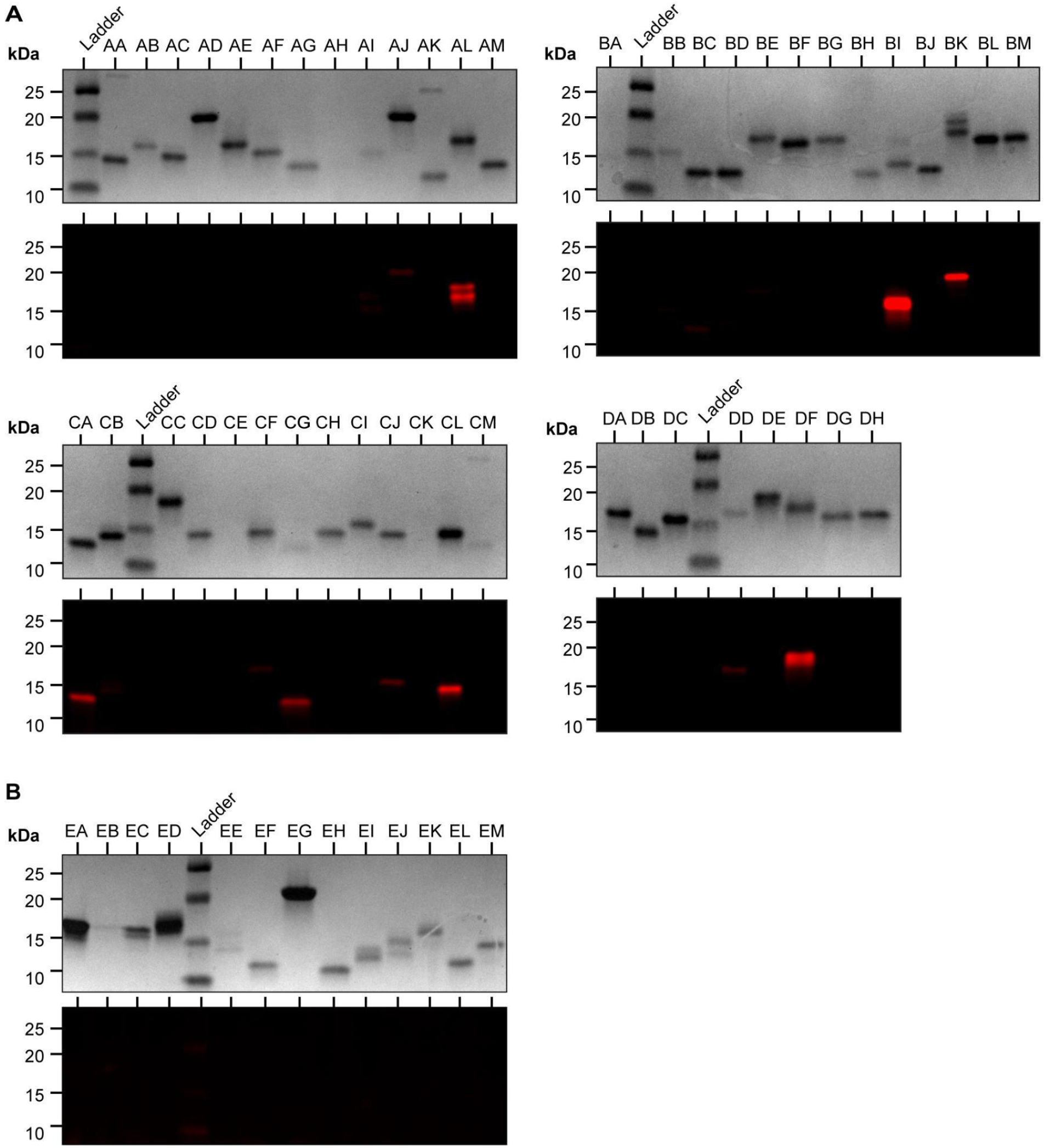
Screening of JF_657_ covalent binder redesigns. **(A)** Coomassie-stained and fluorescent protein SDS-PAGE gel imaging was performed to screen for potential covalent JF_657_ binder. Each design was incubated with excess JF_657_ (10 µM) at 37 °C for 12 h, prior to loading on the protein gel. The co-localization of the far-red fluorescent signal with the protein band indicates the formation of a covalent adduct. Twelve designs showing co-localization of the far-red fluorescent band with the protein band (AJ, AL, BI, BK, BD, CA, CB, CG, CJ, CL, DD, DF) were identified as potential covalent binders. The sample in lane BI, which exhibited the highest fluorescence intensity, was selected for further characterization (**Fig. S16**) and subsequently reported as NovoTag_657_^cv^. **(B)** Coomassie-stained and fluorescent protein SDS-PAGE gel imaging of non-covalent JF_657_ binders, subjected to identical incubation conditions as (A), showed no far-red fluorescent bands co-localizing to coomassie-stained protein bands. This control demonstrated that covalent bond formation between the protein and dye is strictly required to retain the fluorescent signal under denaturing conditions.

**Fig. S16.**
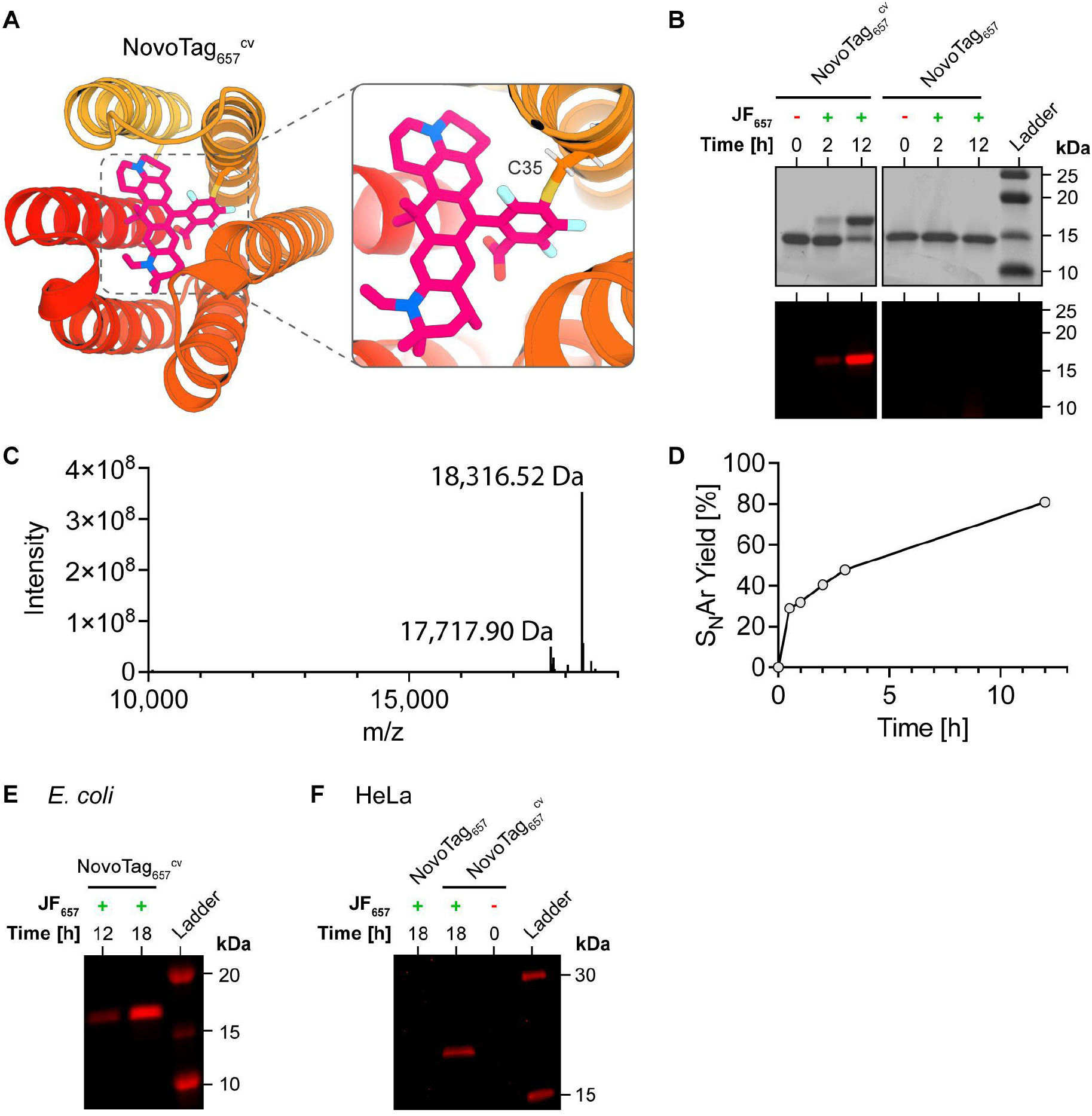
Covalent dye binding characterization of NovoTag_657_^cv^. **(A)** Design model of NovoTag_657_^cv^ with a catalytic cysteine. **(B)** SDS-PAGE of the selected covalent design NovoTag_657_^cv^ (Lane BI in Fig. S15) and the non-covalent NovoTag_657_. For each sample, 5 µM binder was incubated with 10 µM JF_657_ for 2 or 12 h, or not treated with JF_657_ as a control. Coomassie-stained (top) and fluorescence (bottom) images of the gel are shown. The co-localization of the far-red fluorescent signal with the protein band indicates the formation of a covalent adduct. The absence of fluorescent bands in the non-covalent design, confirms the specificity of the covalent bond formation. **(C)** Deconvoluted mass spec trace of NovoTag_657_^cv^ after a 12 h incubation with 50 µM JF_657_, showing peaks corresponding to both labeled and unlabeled species: [NovoTag_657_^cv^ + JF_657_ + H-Met-H-F]^+^ showed an observed peak of 18,316.52 Da, with an expected peak at 18,316.6 Da. [NovoTag_657_^cv^ + H-Met]^+^ showed an observed peak of 17,717.90 Da, with an expected peak at 17,717.87 Da. **(D)** Labeling kinetics of NovoTag_657_^cv^ (10 μM protein incubated with 50 μM JF_657_) were monitored via mass spectrometry over a 12 h incubation period at 37 °C, with integrated area of deconvoluted mass peaks used to quantify the labeling yield. Data represent the mean of three replicates. **(E)** SDS-PAGE of *E. coli* BL21 whole-cell lysate indicating covalent labeling of NovoTag_657_^cv^. Cells were incubated with 1 µM JF_657_ for 12 or 18 h. **(F)** SDS-PAGE of HeLa whole-cell lysate indicating covalent labeling of NovoTag_657_^cv^. As a control, non-covalent NovoTag_657_ was used. Cells were incubated with 1 µM JF_657_ for 18 h.

**Fig. S17.**
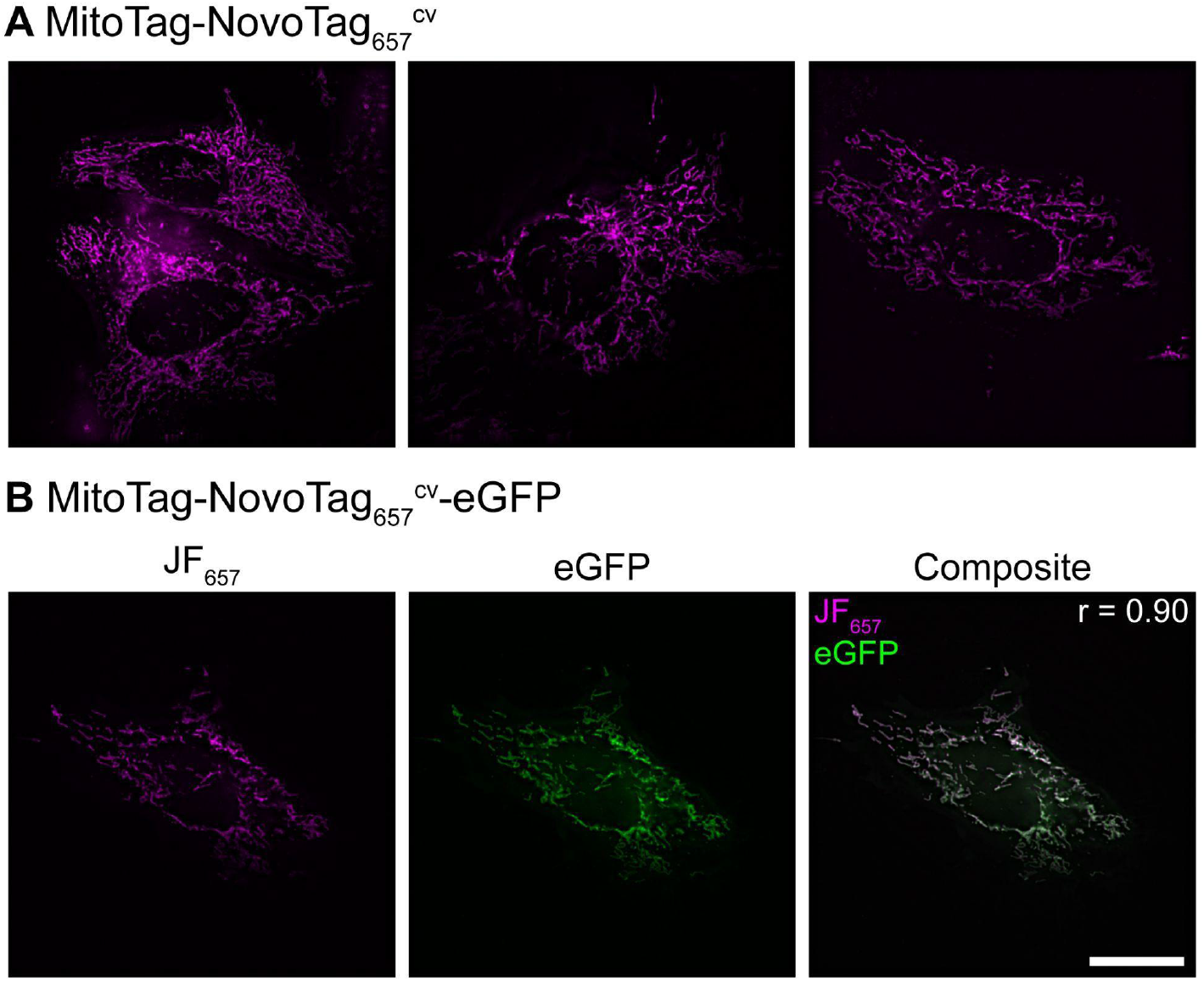
Fluorescence microscopy of covalent NovoTag_657_^cv^ in live cells. **(A)** HeLa cells were transfected with MitoTag-NovoTag_657_^cv^. Cells were incubated with 5 nM of the JF_657_ for 15 min. After three washing steps, live cells were imaged on an OMX SR microscope. Three individual cells are shown. **(B)** HeLa cells were transfected with MitoTag-NovoTag_657_^cv^-eGFP. Cells were incubated with 5 nM of the JF_657_ for 15 min. After three washing steps, live cells were imaged on an OMX SR microscope. Individual channels are shown for JF657 (magenta) and eGPF (green). In addition, a composite image is shown. Pearson’s correlation coefficient (r) is shown for the composite. Scale bar: 20 µm.

**Fig. S18.**
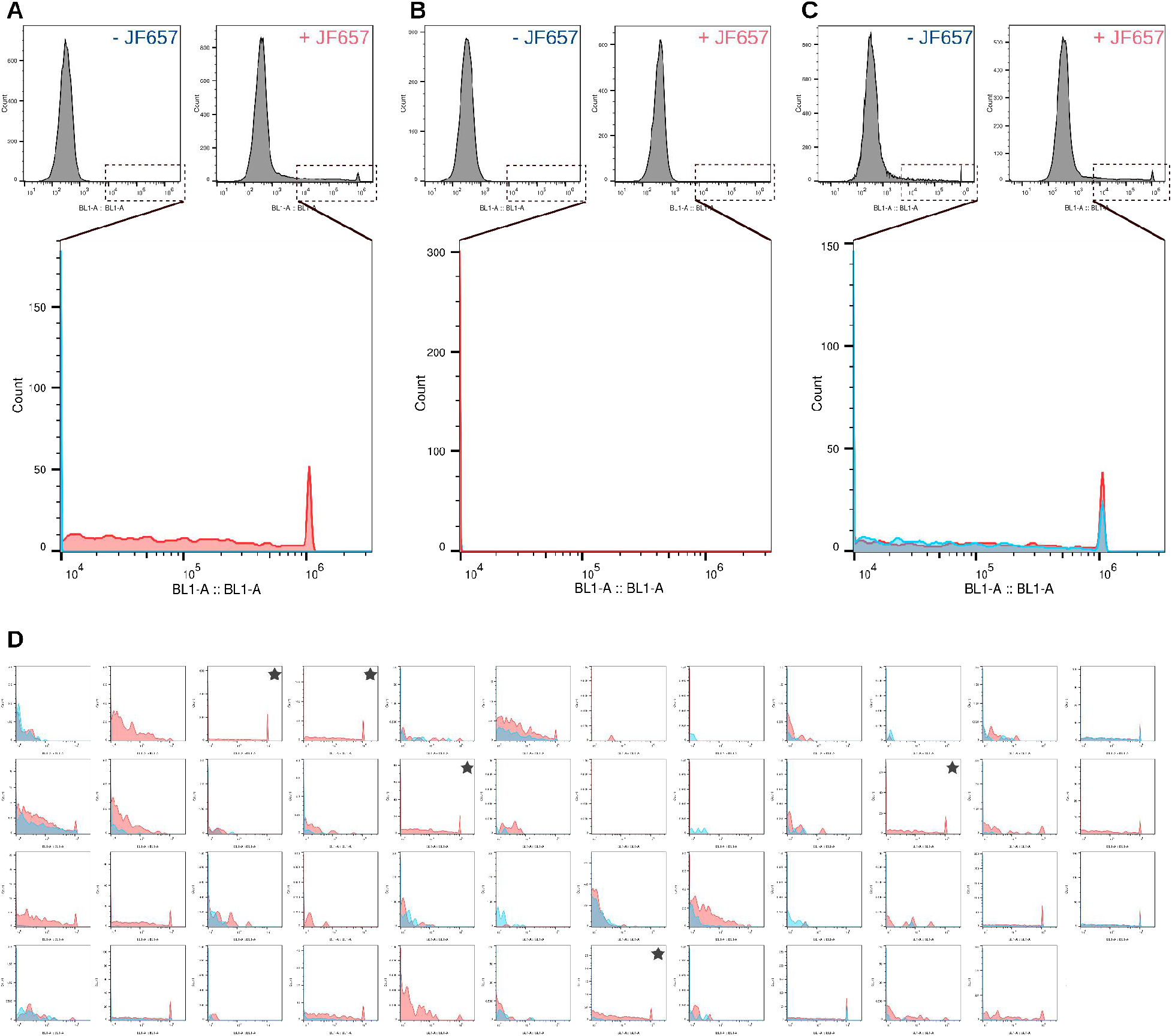
Mammalian two-hybrid screen for JF_657_ split binders. **(A – D)** Flow cytometry histograms of mNeonGreen expression in HEK293T triple transfected with designed JF_657_ split-binder versions and a reporter plasmid encoding for mNeonGreen. Plots represent cells 24 h post JF_657_ incubation (red), and untreated controls (blue), with x-axis representing green fluorescence intensity on log scale (laser excited at 488 nm). mNeonGreen expression indicates designed protein-protein interaction from the split binders. Exemplary histograms are provided for the following design variants: split design with induced mNeonGreen expression upon JF_657_ treatment indicating controlled dimerization (A), split design that does not induced mNeonGreen expression upon JF_657_ treatment (B), and a constitutive (non-specific) dimer variant (C). (**D**) Flow cytometry histograms of all 47 designs screened. Designs indicated by asterisk (similar to profile shown in A) were selected for further screening by colocalization analysis (**Fig. S19**).

**Fig. S19.**
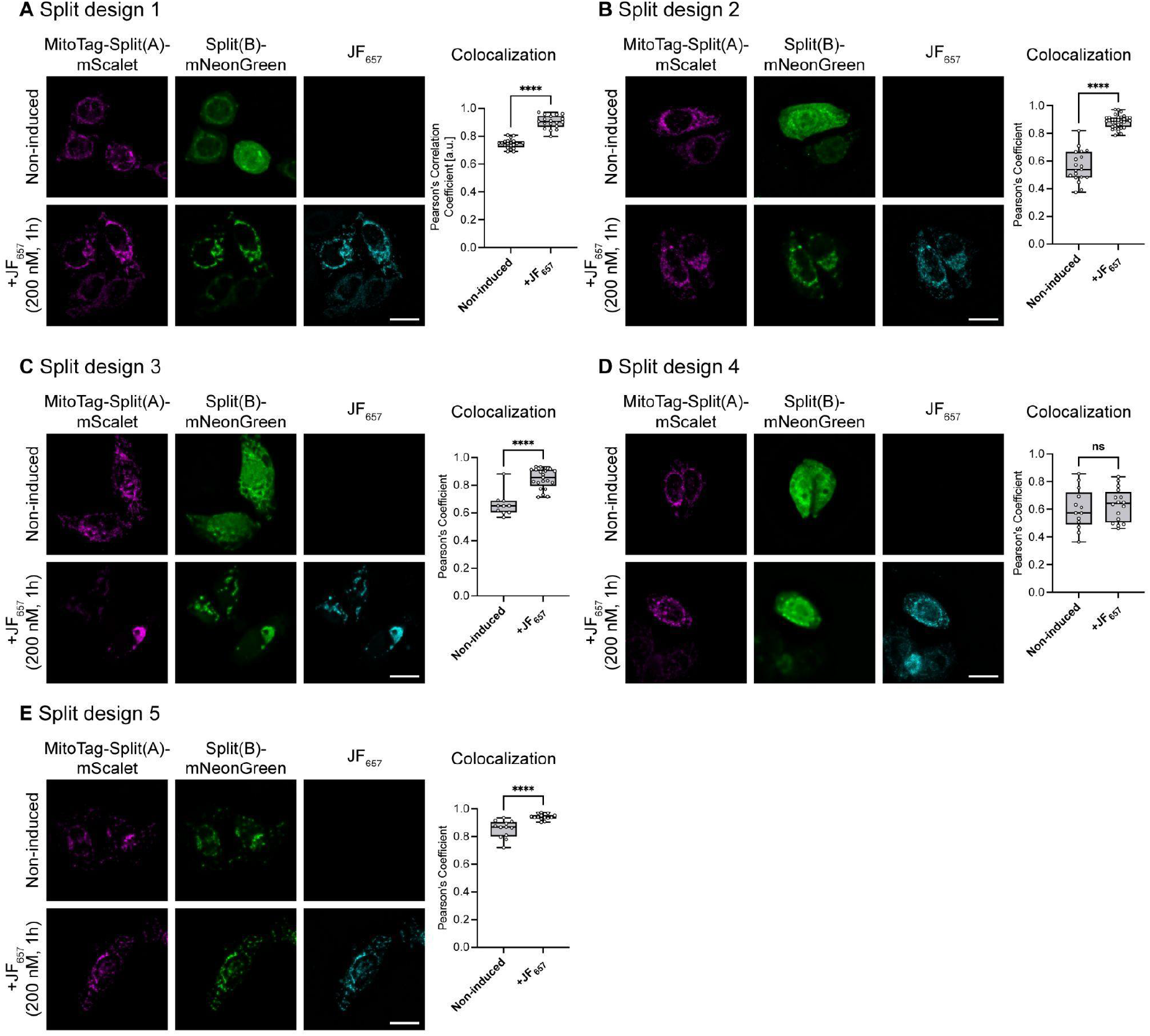
Colocalization screen for JF_657_ split binders. **(A – E)** Five different split binder variants identified from the mammalian two-hybrid screen (Fig. S18) were further screened by fluorescence microscopy. HeLa cells were transfected with MitoTag-split(A)-mScarlet-P2A-split(B)-mNeonGreen and incubated with 200 nM JF_657_ for 1 h. After three washing steps, cells were chemically fixed, and fluorescence widefield images were acquired on a Nikon Ti-Eclipse. The exemplary images show a maximum intensity projection of a Z-stack. Images were deconvolved with Huygens Professional (SVI). Colocalization was analyzed by calculating the Pearson’s correlation coefficient between mScarlet and mNeonGreen. The split design in panel B was selected as the best design. For each sample, between 10 and 34 individual cells were analyzed. All data are shown as Box and Whiskers plots indicating the median (center of box), 25% and 75% quartiles (bounds of box), minimum and maximum values (bars). Each data point represents one cell. Unpaired two-sided Welch’s t-tests were conducted. Statistical significance: ns (non significant) p > 0.05, ∗∗∗∗ p < 0.0001. Scale bar: 20 µm.

**Fig. S20.**
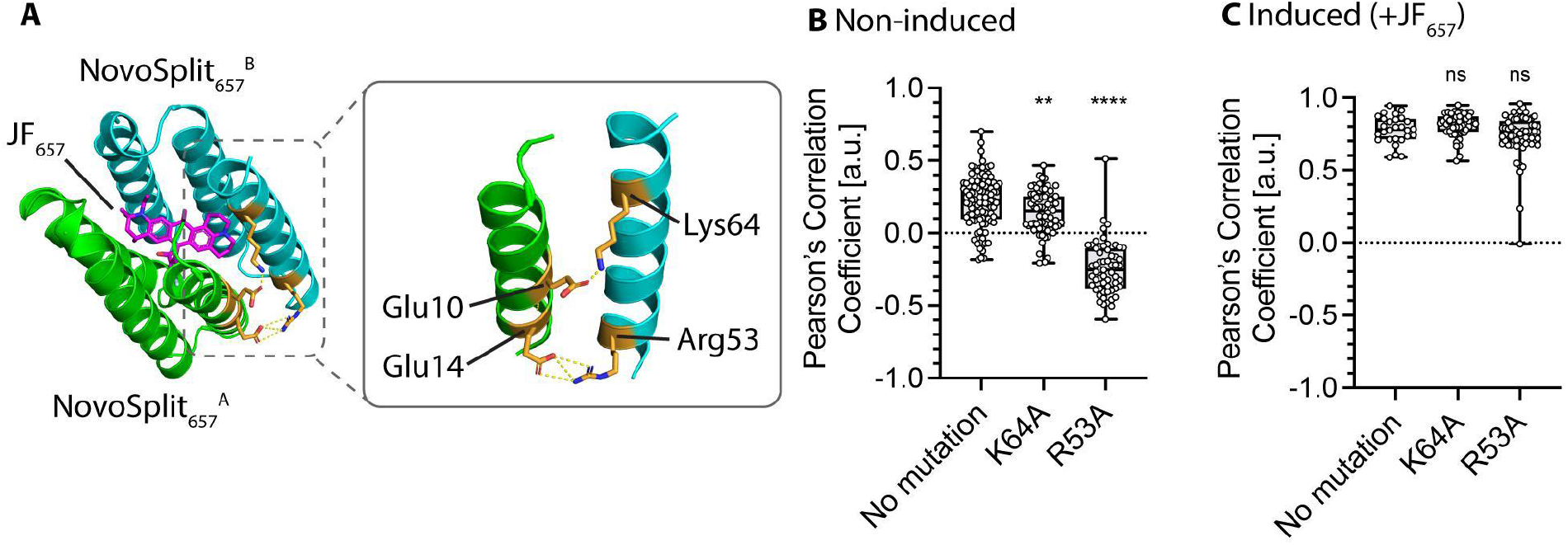
Structure-guided site-directed mutagenesis of NovoSplit_657_ to prevent non-specific self-dimerization. **(A)** Based on the AF2 model of NovoSplit_657_, two amino acid pairs, Glu10–Lys64 and Glu14–Arg53, between NovoSplit_657_^A^ and NovoSplit_657_^B^, were identified as potentially forming salt bridges (yellow dotted lines). We selected Lys64 and Arg53 as potential residues whose mutants to Ala should eliminate the formation of these interactions between the two monomers. **(B & C)** HeLa cells were transfected with MitoTag-NovoSplit_657_^A^-mStayGold and NovoSplit_657_^B^-mCherry, either without any mutation or with Lys64Ala or Arg53Ala mutation in NovoSplit_657_^B^. Colocalization analysis between NovoSplit_657_^A^ and NovoSplit_657_^B^ in the non-induced state (B) revealed a reduced correlation coefficient for both mutations, with r = 0.14 ± 0.14 for Lys64Ala, and r = -0.24 ± 0.18 for Arg53Ala. Induction of dimerization with JF_657_ (200 nM for 1 h) was not impaired by either mutation (C). Data is shown as Box and Whiskers plots, indicating the median (center of the box), 25% and 75% quartiles (box bounds), and minimum and maximum values (bars). Each data point represents one cell. Unpaired two-sided Welch’s t-tests were conducted to compare each mutation to the non-mutated control. Statistical significance: ns (non significant) p > 0.05, ∗∗ p < 0.01, and ∗∗∗∗ p < 0.0001.

**Fig. S21.**
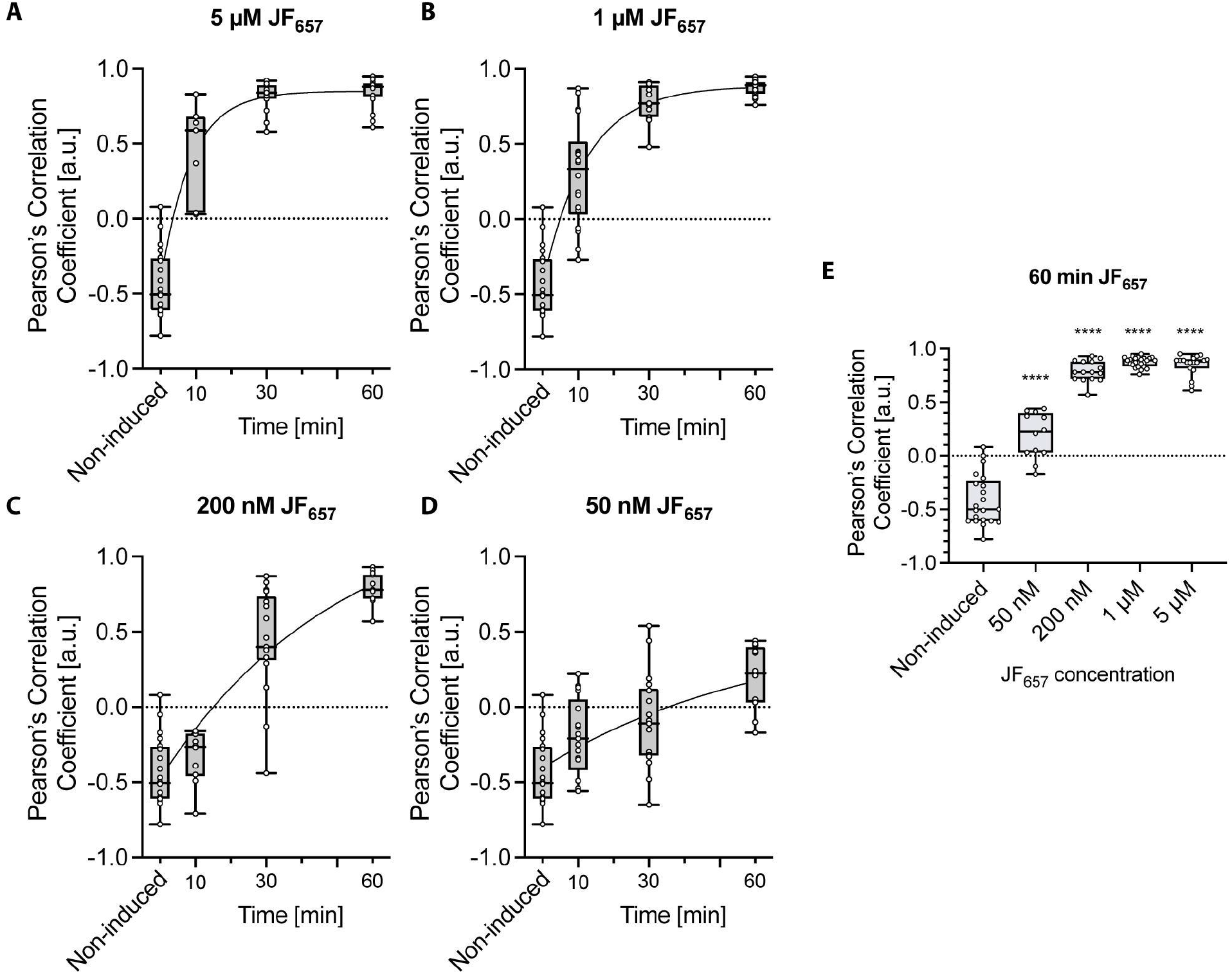
NovoSplit_657_ binding kinetics in HeLa cells. (**A – D**) The NovoSplit system (see schematic in Fig. 4C) was transfected into HeLa cells and treated with different concentrations of JF_657_: 5 µM (A), 1 µM (B), 200 nM (C), and 50 nM (D). Cells were fixed at specific time points post JF_657_ treatment (10 min, 30 min, and 60 min), fluorescent images were acquired, and Pearson’s correlation coefficients between MitoTag-mStayGold-NovoSplit_657_^A^ and mCherry-NovoSplit_657_B were determined. A nonlinear regression (one-phase association) was fitted, and the half-times (t_1/2_) were determined: t_1/2_ = 6 min for 5 µM JF_657_, t_1/2_ = 9 min for 1 mM JF_657_, t_1/2_ = 35 min for 200 nM JF_657_, and t_1/2_ = 44 min for 50 nM JF_657_. **(E)** Comparison of MitoTag-mStayGold-NovoSplit_657_^A^ and mCherry-NovoSplit_657_B colocalization with different JF_657_ concentrations after 60 min of JF_657_ treatment. Unpaired two-sided Welch’s t-tests were conducted to compare each condition to the non-induced control. Statistical significance: ∗∗∗∗ p < 0.0001. All data are shown as Box and Whiskers plots, indicating the median (center of the box), 25% and 75% quartiles (box bounds), and minimum and maximum values (bars). Each datapoint represents one cell.

**Fig. S22.**
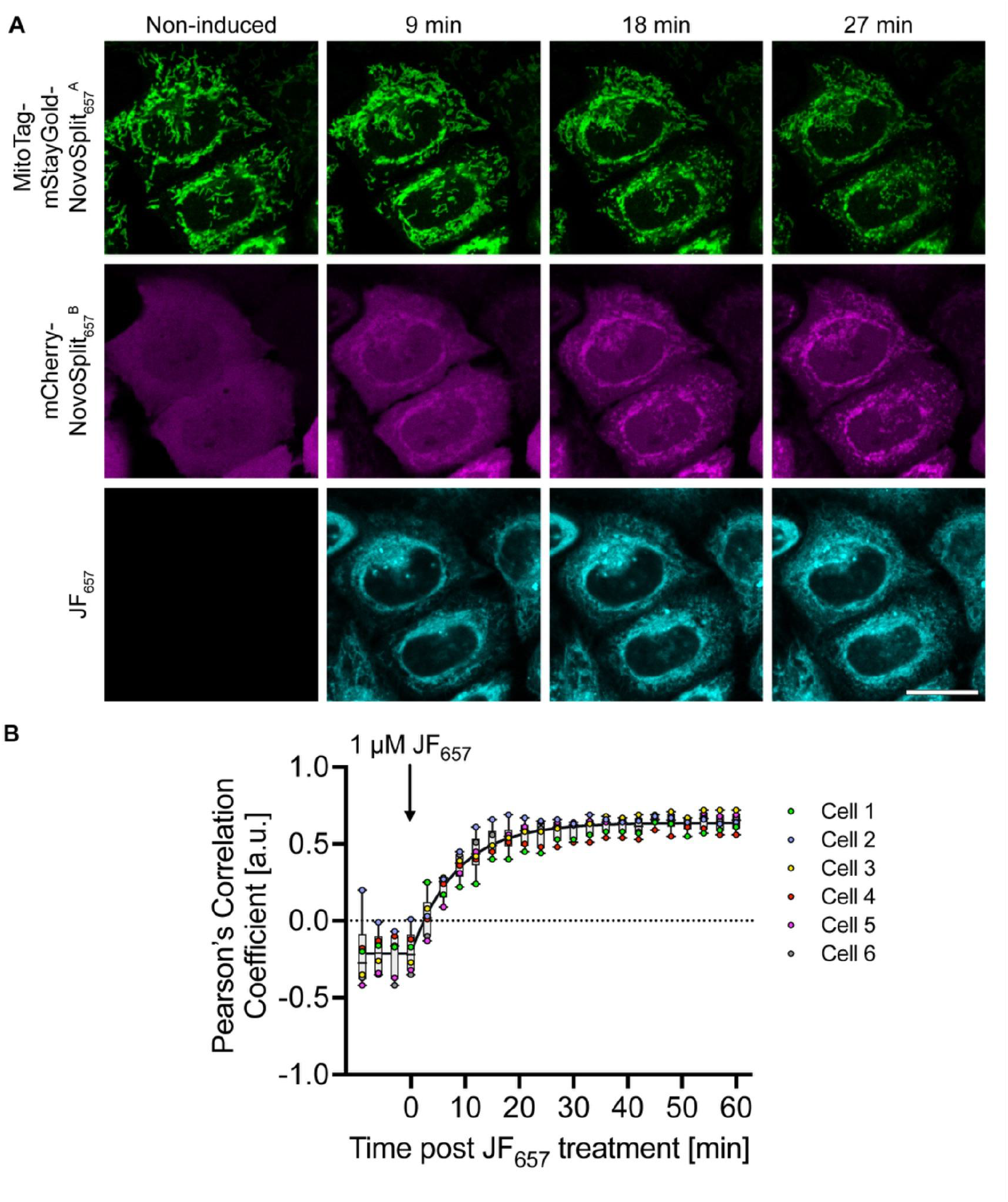
Live cell imaging of NovoSplit_657_ induced dimerization in HeLa cells. **(A)** The NovoSplit system (see schematic in Fig. 4C) was transfected in HeLa cells and live cell fluorescence imaging was performed. Every 3 min, a frame was acquired. Non-induced state and representative time points after 1 µM JF_657_ treatment (9, 18, 27 min) are shown. MitoTag-mStayGold-NovoSplit_657_^A^ is shown in green, mCherry-NovoSplit_657_^B^ in magenta, and JF_657_ in cyan. **(B)** For each time point, the Pearson’s correlation coefficients between MitoTag-mStayGold-NovoSplit_657_^A^ and mCherry-NovoSplit_657_^B^ were determined. Analysis was performed for 6 cells. A nonlinear regression (plateau followed by one phase association) was fitted to the mean of the 6 analyzed cells, and a half-time of t_1/2_ = 5.8 min (95% confidence interval (CI): 5.0 – 6.7 min) was determined. Scale bar: (A) 20 µm.

**Fig. S23.**
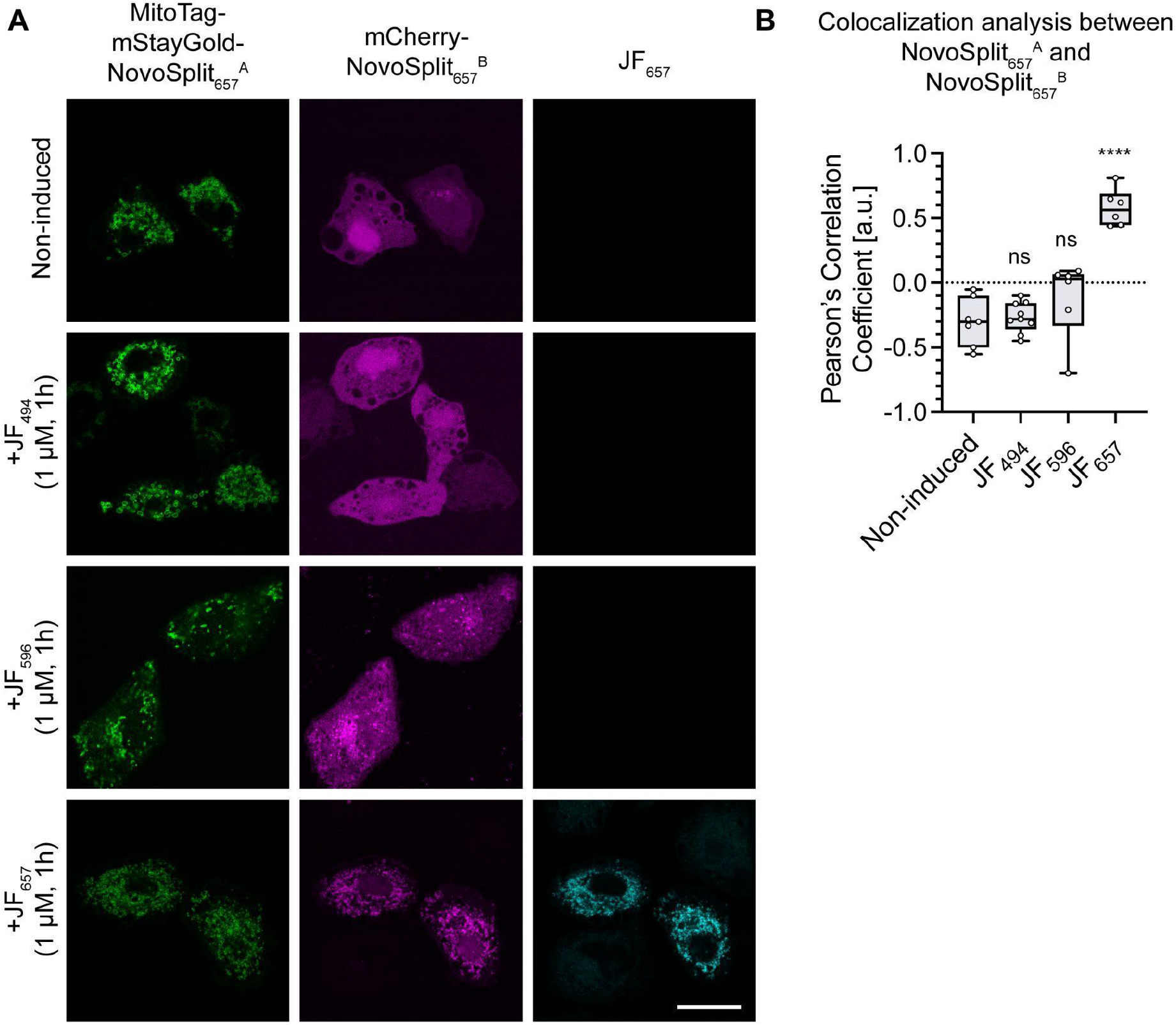
Analysis of NovoSplit_657_ dye binding specificity in HeLa cells. **(A)** The NovoSplit system (see schematic in Fig. 4C) was transfected in HeLa cells and cells were treated with 1 µM of one of the three JF dyes (JF_657_, JF_596_, or JF_494_) for 1 h. Cells were fixed, washed, and fluorescent images acquired on a Zeiss AiryScan 980 confocal microscope. **(B)** Pearson’s correlation coefficients between MitoTag-mStayGold-NovoSplit_657_^A^ and mCherry-NovoSplit_657_^B^ were determined. Unpaired two-sided Welch’s t-tests were conducted to compare each condition to the non-induced control. Statistical significance: ns (non significant) p > 0.05, ∗∗∗∗ p < 0.0001. All data are shown as Box and Whiskers plots, indicating the median (center of the box), 25% and 75% quartiles (box bounds), and minimum and maximum values (bars). Each datapoint represents one cell. For each condition, 6-9 individual cells were analyzed.

## Supplementary Tables

**Table S1.** Crystal structure data collection and refinement statistics for NovoTag_657_-apo and -holo. Single crystal used for each data collection. Values in parentheses are for the highest-resolution shell.

|  | **NovoTag<sub>657</sub>-apo**<br>**(PDB Code: 9PVL)** | **NovoTag<sub>657</sub>-holo**<br>**(PDB Code: 11MQ)** |
| --- | --- | --- |
| **Cell parameters** |  |  |
| Space group | *P 2<sub>1</sub>* | *P 2<sub>1</sub> 2<sub>1</sub> 2<sub>1</sub>* |
| a,b,c (Å) | 33.36, 123.99, 56.64 | 32.29, 56.63, 67.49 |
| α, β, γ (°) | 90, 92.28, 90 | 90, 90, 90 |
| Resolution (Å) | 33.33 - 1.44 (1.51 - 1.44)* | 67.49 - 2.37 (2.46 - 2.37) |
| *R<sub>merge</sub>* | 0.076 (0.897) | 0.588 (1.361) |
| *R<sub>pim</sub>* | 0.035 (0.430) | 0.187 (0.440) |
| *I*/*σ*(*I*) | 10.9 (1.9) | 6.0 (1.9) |
| *CC*<sub>1/2</sub> | 0.998 (0.625) | 0.962 (0.360) |
| Completeness (%) | 100 (100) | 100 (100) |
| Redundancy | 6.6 (6.1) | 11.9 (11.2) |
| **Refinement** |  |  |
| Resolution (Å) | 33.33 - 1.44 (1.47 - 1.44) | 43.38 - 2.38 (2.62 - 2.38) |
| No. reflections | 83269 | 5319 |
| *R<sub>work</sub>* / *R<sub>free</sub>* | 0.2112 (0.3099)/<br>0.2447 (0.2970) | 0.2455 (0.2931)/<br>0.2904 (0.3097) |
| No. of atoms |  |  |
| Protein | 4432 | 1108 |
| Ligand | n/a | 58 |
| Solvent | 392 | 9 |
| Ramachandran favored/allowed/ outlier (%) | 99.81/ 019/ 0.00 | 97.74/ 2.26/ 0.00 |
| RMSD |  |  |
| Bond lengths (Å) | 0.008 | 0.002 |
| Bond angles (°) | 1.027 | 0.603 |
| *B<sub>factors</sub>* (Å<sup>2</sup>) |  |  |
| Protein | 23 | 35 |
| Ligand |  | 39 |
| Solvent | 29 | 39 |

**Table S2.**
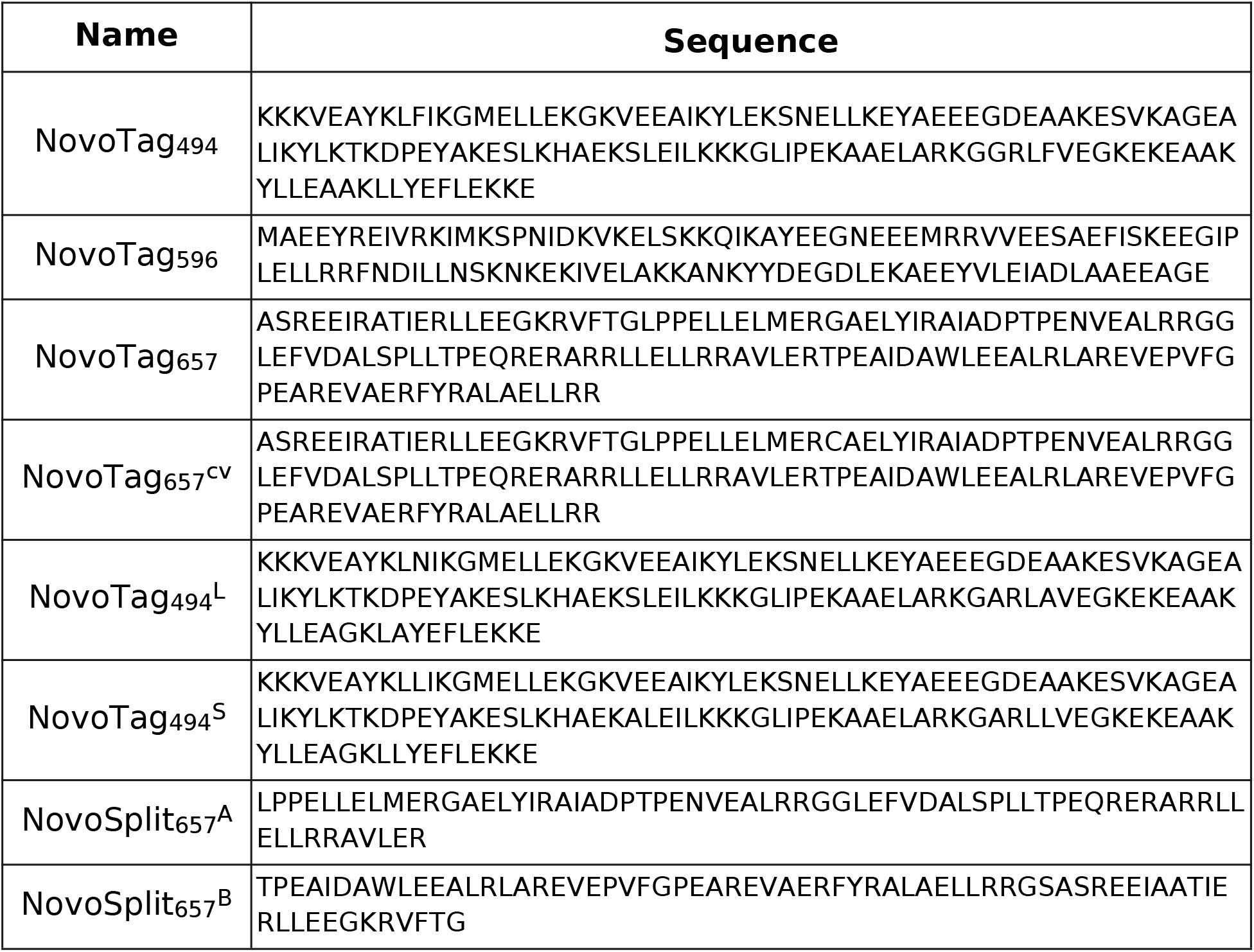
Amino acid sequences of the NovoTag and NovoSplit systems generated in this work.

**Table S3.** Plasmids generated in the study.

| **Plasmid Name** | **Addgene ID** |
| --- | --- |
| 2×FYVE-NovoTag<sub>494</sub> | 255873 |
| Tom70-NovoTag<sub>596</sub> | 255874 |
| NovoTag<sub>596</sub>-Emerin | 255875 |
| Tom70-NovoTag<sub>657</sub> | 255876 |
| H2B-NovoTag<sub>657</sub> | 255877 |
| Tom70-NovoTag<sub>494</sub><sup>S</sup> | 255878 |
| H2B-NovoTag<sub>494</sub><sup>L</sup> | 255879 |
| Tom70-NovoTag<sub>657</sub><sup>CV</sup> | 255880 |
| Tom20-mStayGold-NovoSplit<sub>657</sub><sup>A</sup> | 255881 |
| mCherry-NovoSplit<sub>657</sub><sup>B</sup> | 255882 |
| Tom70-LHD(A)-NovoSplit<sub>657</sub><sup>A</sup>-<br>mScarlet | 255883 |
| LHD(B)-NovoSplit<sub>657</sub><sup>B</sup>-mNeonGreen | 255884 |
| NovoSplit<sub>657</sub><sup>B</sup>-mNeonGreen | 255885 |

## Supplementary Movies

**Movie S1: Photostability comparison of JF_657_:NovoTag_657_ and JFX_650_-HALO:HaloTag7.** Movie related to **Fig. 2C** and **Fig. S10**. Scale bar: 20 µm

**Movie S2: Live cell fluorescence imaging of NovoSplit_657_ in HeLa cells.** Movie related to **Fig. S22**. Scale bar: 30 µm.

**Movie S3: Design pipeline trajectory.** Movie related to design pipeline trajectory of NovoTag_494_. Spheres represent Cα. Ligand and key interacting residues were represented in sticks.

## Notes

### Competing Interest Statement

This work is filed for a provisional patent.

### Summary of Updates

change distribution to cc. change distribution to cc

